# The impact of ribosomal interference, codon usage, and exit tunnel interactions on translation elongation rate variation

**DOI:** 10.1101/090837

**Authors:** Khanh Dao Duc, Yun S. Song

**Affiliations:** Computer Science Division, University of California Berkeley, Berkeley, CA 94720; Department of Statistics, University of California Berkeley, Berkeley, CA 94720; Chan Zuckerberg Biohub, San Francisco, CA 94158, USA

## Abstract

Previous studies have shown that translation elongation is regulated by multiple factors, but the observed heterogeneity remains only partially explained. To dissect quantitatively the different determinants of elongation speed, we use probabilistic modeling to estimate initiation and local elongation rates from ribosome profiling data. This model-based approach allows us to quantify the extent of interference between ribosomes on the same transcript. We show that neither interference nor the distribution of slow codons is sufficient to explain the observed heterogeneity. Instead, we find that electrostatic interactions between the ribosomal exit tunnel and specific parts of the nascent polypeptide govern the elongation rate variation as the polypeptide makes its initial pass through the tunnel. Once the N-terminus has escaped the tunnel, the hydropathy of the nascent polypeptide within the ribosome plays a major role in modulating the speed. We show that our results are consistent with the biophysical properties of the tunnel.

## INTRODUCTION

Ribosome profiling [1–3] is a powerful transcriptome-wide experimental protocol that utilizes high-throughput sequencing technology to provide detailed positional information of ribosomes on translated mRNA transcripts. As a useful tool to probe post-transcriptional regulations of gene expression, ribosome profiling has notably been used to identify translated sequences within transcriptomes, to monitor the process of translation and the maturation of nascent polypeptides *in vivo,* and to study limiting determinants of protein synthesis (see recent reviews [4–6] for an overview of diverse applications of the technique). In addition, since the ribosome occupancy at a given position reflects the relative duration of time spent at that position, ribosome profiling provides an unprecedented opportunity to study the local translational dynamics [7]. However, the precise relation between the observed footprint densities and the corresponding translation elongation rates remains elusive [6], thus making it difficult to interpret ribosome profiling data.

One factor that may affect the translation elongation speed is ribosomal interference, which occurs when slow translocation of a ribosome at a certain site blocks another one preceding it (an extreme case being ribosomal pausing, occurring both in eukaryotes and bacteria [8]). Because the information provided by ribosome profiling is marginal probability density (in the sense that it does not capture the joint occupancy probability of multiple ribosomes on the same transcript), it is not possible to observe ribosomal interference directly from data and therefore quantifying the role of interference in limiting the elongation speed has remained challenging. Furthermore, a further challenge for inference arises from the potential omission of stacked ribosomes (i.e., multiple ribosomes that are less than a few codons apart) in the current ribosome profiling protocol [9–12]. In most studies, positional distributions of ribosomes along the open reading frame (ORF) are inferred from protected mRNA fragments of lengths 27-31 nt that presumably reflect the size of the 60S ribosomal subunit (28-29 nt in *S. cerevisiae* or 30-31 nt in mammalian cells). However, gradient footprint profile has detected longer protected fragments of 40-65 nt which can be attributed to two closely stacked ribosomes that accumulate when the leading ribosome is stalled [11,13]. Not taking these longer mRNA fragments into account in the ribosome profile may thus produce biased estimates of ribosome densities, and, as a consequence, of elongation rates.

Over the past few years, multiple studies have tried to utilize ribosome profiling data to identify the key determinants of the protein production and translation rates, but have arrived at contradictory results [14–22]. Due to the vast complexity of the different biophysical mechanisms involved in the decoding and translocation of the ribosome along the mRNA, it is indeed a challenging problem to disentangle the composite factors that can modulate the elongation speed for a given transcript sequence. Several studies have shown that elongation speed is locally regulated by multiple factors, including tRNA availability and decoding time [21,23,24], mRNA secondary structure [25], peptide bond formation at the P-site [26], and the presence of specific amino acid residues [17,18,27,28] in the nascent polypeptide that interact with the ribosomal exit tunnel [29]. However, the observed heterogeneity in elongation rates along the transcript, notably the so-called 5′ “translational ramp” [1], remains only partially explained [16,30].

Here, we provide new insights into the major determinants of the translation dynamics, by identifying features that can explain a large portion of the variation in the mean elongation rate along the transcript, particularly the 5′ translational ramp. **W**e also present a new statistical method that can be used to obtain accurate estimates of initiation and local elongation rates from ribosome profiling and RNA-seq data. Our approach is based on a probabilistic model that takes into account the principal features of the translation dynamics, and it allows us to quantify the extent of ribosomal interference (not directly observable from data) along the transcript.

## RESULTS

### Estimation of initiation rates and local elongation rates

We developed an inference procedure based on an extended version of the biophysical TASEP model [31] (see Figure 1, **Materials and Methods**, and Sections 1-3 of **Supporting Information**) to estimate transcript-specific initiation and local elongation rates from ribosome profiling and RNA-seq data. In our model, the initiation rate is the exponential rate at which the P-site of a lower ribosomal subunit arrives at the start codon and the upper ribosomal subunit gets assembled, while the elongation rate at a given codon position is the rate at which the A-site of a ribosome occupying that position translocates to the next downstream codon. Here, both events are conditioned on there being no other ribosomes in front obstructing the movement.

**Figure 1.**
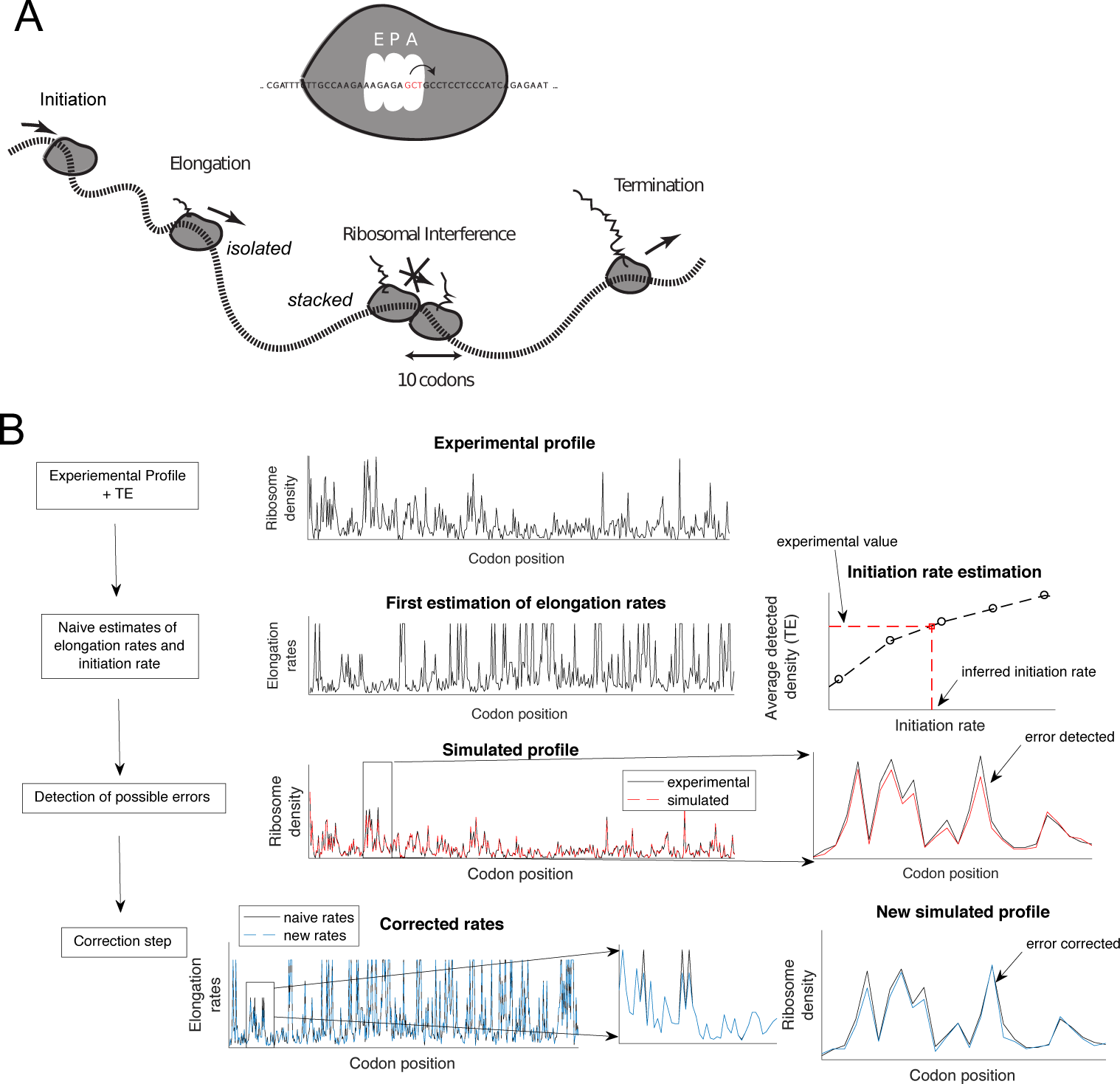
Illustration of translation dynamics and inference from experimental data. **A.** A representation of the mathematical model of translation. Initiation corresponds to an event where the A-site of a ribosome enters the second codon position, while elongation corresponds to a movement of the ribosome such that its A-site moves to the next downstream codon. Both events are conditioned on there being no other ribosomes in front obstructing the movement. The ribosome eventually reaches a stop codon and subsequently unbinds from the transcript. In our main simulations, we say that a ribosome is undetected when the distance between the A-sites of consecutive ribosomes is ≤ 12 codons. **B.** A schematic description of our inference procedure. Given a ribosome profile and a measure of average density (TE), we first approximate the position-specific elongation rates by taking the inverse of the observed footprint number. Then, we use simulation to search over the initiation rate that minimizes the difference between the experimental density and the one obtained from simulation. **W**e then iteratively refine these estimates: **W**e compare the simulation result with the experimental ribosome profile and detect “error-sites” where the absolute density difference is larger than a chosen threshold. If error-sites are found, we start with the one closest to the 5′-end, and jointly optimize the initiation rate and the elongation rates in a neighborhood of this error-site to minimize the error between the simulated and observed profiles. Using these new parameters, we then re-detect possible error-sites located downstream and repeat the procedure (more details in **Material and Methods**).

For the main part of our analysis, we used flash-freeze ribosome profiling and RNA-seq data of *S. cerevisiae* generated by **W**einberg *et al.* [16]. These data have been shown to have substantial improvements over previous datasets, alleviating protocol-specific biases [16] that can influence interpretation of ribosome-profiling experiments [12]. We ran our inference method on a subset of 850 genes selected based on length and footprint coverage (see **Materials and Methods**), and tested its accuracy (detailed in Figure S1 and Figure S2). Of these, 383 genes (45%) did not require any corrections after the first step of our inference procedure, which means that only the initiation rate was fitted to match the data (the inverse of the observed density was used to estimate the elongation rate, see **Materials and Methods**). For the remaining 467 genes, the number of sites that required a correction procedure was on average 1.57 per gene (std = 0.925) (Figure S2A). A more detailed analysis and comparison of these two subsets of genes are provided in Section 4 of Supporting Information and Figures S3, S4, and S5, which show that sites with larger relative footprint densities and codons with slowest average elongation rates are more likely to require corrections. Figure 1B is an example illustrating the excellent agreement between the actual ribosome footprint distribution for a specific gene from experiment and the distribution of detected ribosomes obtained from simulation under the extended TASEP model with our inferred initiation and elongation rates. Other comparisons of experimental and simulated profiles are provided in Figure S6. As shown below, the results from running our method on other ribosome profiling datasets [19,32] (see Section 5 of Supporting Information) are consistent with our results on Weinberg *et al.’s* data.

### Inference of possible omissions of stacked ribosomes from ribosome profiles

The standard ribosome profiling protocol selects for isolated ribosomes occupying 27 and 31 nt, so longer mRNA fragments protected by closely-stacked ribosomes (separated by ≤ 2 codons) are possibly not included in the experimental data, thus making the ribosome footprint distribution inaccurate in regions of high traffic [9–13]. To infer the extent of such omissions in **W**einberg *et al.’s* experimental ribosome profiling data, we compared the translation efficiency (TE) to the measurement of per-gene ribosome density from polysome profiling carried out by Arava *et al.* [33] (for 588 genes common to both datasets). TE is the ratio of the RPKM measurement for ribosomal footprint to the RPKM measurement for mRNA [1], where RPKM corresponds to the number of mapped reads per length of transcript in kilo base per million mapped reads. In other words, it quantifies for each gene the average number of detected ribosomes per single transcript, up to a normalization constant. When the total ribosome density of a gene is low, it coincides with the TE. We therefore determined the normalization constant (0.83) by linearly fitting the TE to the total ribosome densities measured by Arava *et al.* for values less than 1 ribosome per 100 codons (see Figure 2A). Interestingly, when we looked at a subset of higher-density transcripts (> 1 ribosome per 100 codons) contained in Arava *et al.’s* dataset, we found that the normalization constant obtained by fitting these higher densities was lower (0.61), as shown in Figure 2B. This suggested that for highly occupied transcripts, the density of ribosome inferred from TE underestimates the actual total ribosomal density. Using Pop *et al.’s* data [19] and performing the same comparisons led to similar results (Figure S7A-B).

**Figure 2.**
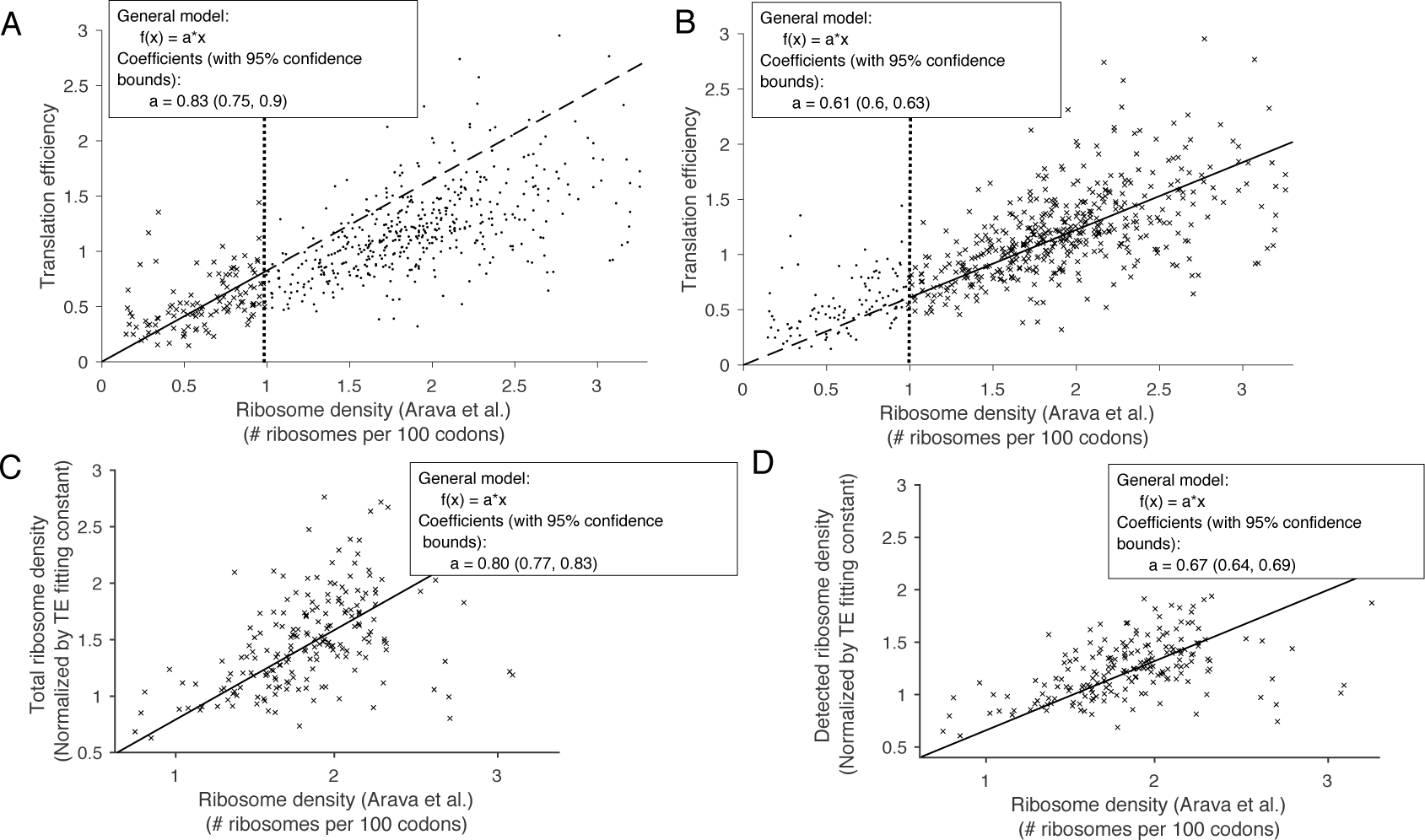
Comparison between translation efficiency (TE) and total ribosome density. All linear fit results are shown in the inset. A. The gene-specific TE for 588 genes from Weinberg *et al*.’ s data [16] (see **Materials and Methods**) against the corresponding total ribosome density (average number of ribosomes per 100 codons) from Arava *et al.* [33]. We performed a linear fit of the points for which the corresponding ribosome density was less than 1 ribosome per 100 codons. **B**. Similar fit as in **A** in the range of ribosome density larger than 1 ribosome per 100 codons. **C**. For the genes (195 in total) that belong to both our main dataset and Arava *et al.*’ s, we compared the simulated total densities obtained using our inferred rates, against the ribosome density from Arava *et al.* **D**. Simulated detected-ribosome densities for the same 195 genes against the ribosome density from Arava *et al.* These results suggest that closely-stacked ribosomes comprise a large fraction of undetected ribosomes, and that our method allows us to correct the TE value to get close to the actual total ribosome density.

To see if our method could accurately capture this difference, we compared the experimentally measured densities (in Arava *et al.)* with our simulated average densities. Specifically, we simulated average densities using the rates inferred from experimental ribosome profiles and TE measurements under three different scenarios of undetected ribosomes (Figure S8): 1) when all closely-stacked ribosomes are detected, 2) when each detection is only partially successful with probability 0.5, and 3) with no detection of the closely-stacked ribosomes. For each scenario, we produced a linear fit of the total ribosome density (obtained by combining detected and undetected ribosomes) against Arava *et al.’s* data. Our goal here was to see which scenario produces a linear-fit coefficient that is the closest to the aforementioned normalization constant (0.83) for low-density transcripts (with < 1 ribosome per 100 codons). The linear-fit coefficient was 0.63 for the first scenario (complete detection of closely-stacked ribosomes), 0.7 for the second (partial detection), and 0.80 for the last (no detection). The last scenario agrees the best with the normalization constant (0.83) for low-density transcripts (Figure 2C). Furthermore, when we used this model to fit the density of only *detected* ribosomes against Arava *et al.’s* data (Figure 2D), we found the normalization constant to be lower (0.67), consistent with the decrease we observed in the fit of the raw TE values for high-density transcripts (Figure 2B). Applying our inference procedure to Pop *et al.’s* data yielded similar results (see Figure S7C-D).

### Inferred rates are consistent with existing results and across different datasets

Upon selecting a model of ribosome profiling with no detection of closely-stacked ribosomes, we used the corresponding estimates to see whether our method could recover what is known in the literature. For the set of genes we considered, we found that the mean time between initiation events varied from 5.5 *s* (5th percentile) to 20 *s* (95th percentile), with median = 10 s. These times are of similar order but shorter than the times found previously [15]. (4 *s* to 233 *s* for the 5th to 95th interpercentile range, with median = 40 s), which can be partially explained by the fact that the set of genes we considered does not include lowly expressed genes (i.e., with low ribosomal density). For the subset of 850 genes that we considered, the corresponding median time between initiation events is indeed substantially lower (by 40%) than that for the whole gene set. In agreement with previous findings [15,16,34], our inferred initiation rates were also positively correlated (Pearson’s correlation coefficient *r* = 0.2646, p-value < 10^−5^) with the 5′-cap folding energy (see **Materials and Methods**) and negatively correlated (r = −0.4, p-value < 10^−5^) with the ORF length (these results are detailed in Figure 3A). We also compared our estimated initiation rates with the ones inferred by Ciandrini *et al.* [34], who develop a simpler approach to infer initiation rates from polysome profile data [35]. The mean initiation times from Ciandrini *et al.* were close to ours (4.6 *s* to 15 *s* for the 5th to 95th interpercentile range, with median = 8 s, Figure S9A), and a direct comparison between the two sets of initiation rates showed a positive correlation (R^2^ = 0.4, see Figure S9B). Applying our method to Pop *et al.’s* dataset (see Figure S10) also leads to a positive correlation (0.31, *p* < 10^−4^), with improved consistency as the sequencing depth increases (see Section 6 of **Supporting Information**).

**Figure 3.**
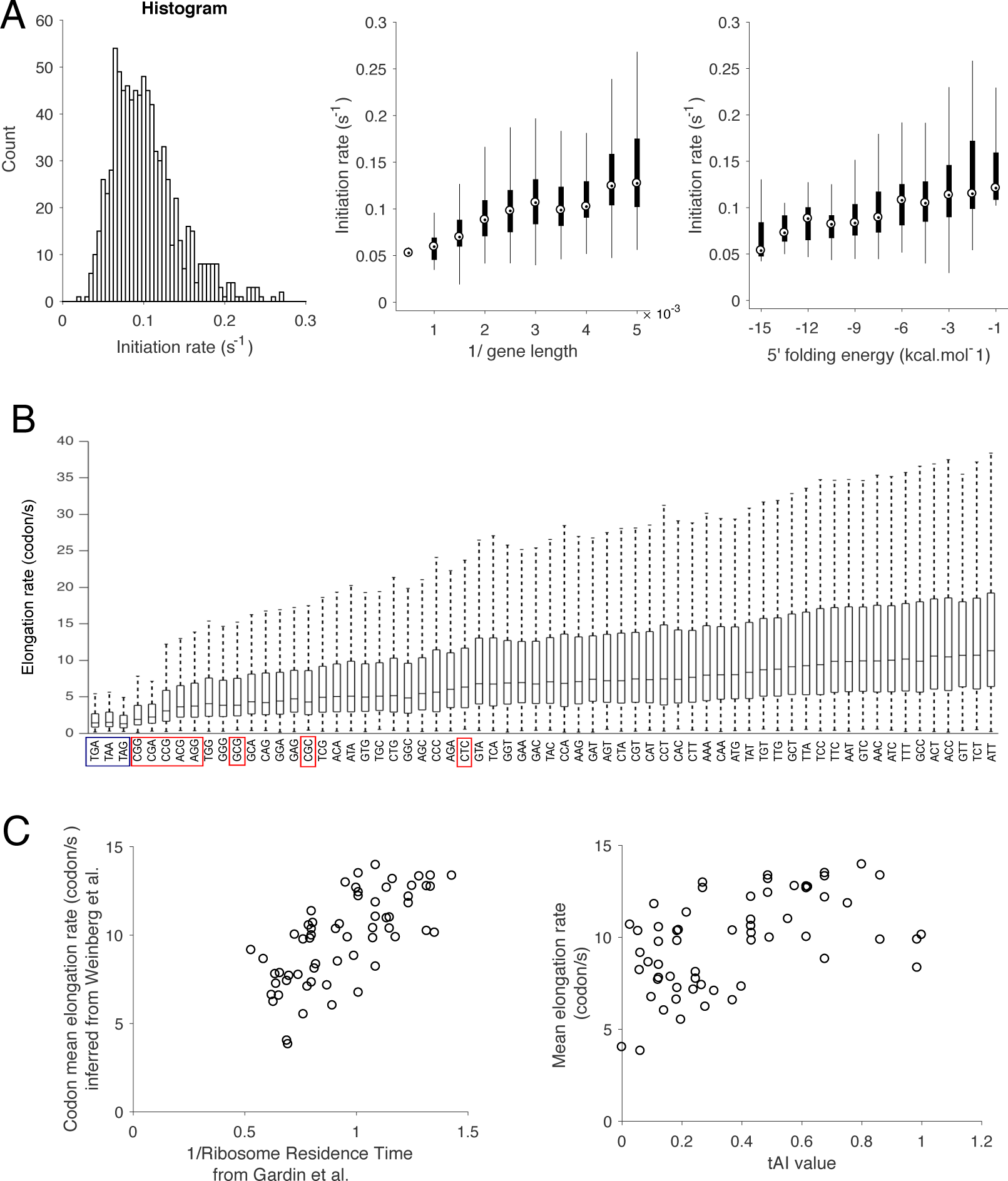
Analysis and comparison of the inferred rates. **A.** (Left) A histogram of inferred initiation rates. (Middle) Comparison between the inferred initiation rates and the inverse of the ORF length of the gene, showing a positive correlation (*r* = 0.44, p-value < 10^−5^, computed for unbinned data). (Right) Comparison between the inferred initiation rates and the 5′-cap folding energy computed in **Weinberg** *et al.* [16], showing a positive correlation (Pearson’s correlation coefficient *r* = 0.2646, p-value < 10^−5^, computed for unbinned data). The interquartile range is indicated by the box, the median by a point inside the box, and upper and lower extreme values by whiskers. **B**. Distribution of codon-specific elongation rates. Stop codons are boxed in blue, while the eight low-usage codons reported by Zhang *et al.* [75] are boxed in red. **C**. Comparison between the codon-specific mean elongation rates computed from B and (Left) the inverse of the codon mean "ribosome residence time” (RRT) estimated by Gardin *et al.* [21], and (Right) the tAI value, computed by Tuller *et al.* [14].

To verify that our method effectively captured the dynamics associated with a specific codon at the A-site, we separated the inferred elongation rates according to their corresponding codon (the resulting distributions are shown in Figure 3B). We observed that codon-specific mean elongation rate (MER) was positively correlated with the inverse of the codon-specific A-site decoding time estimated from Gardin *et al.* [21] *(r =* 0.7, p-value < 10^−5^, see Figure 3C), supporting that different codons are decoded at different rates at the A-site. We then compared these MER with the ones estimated by applying our method to another flash-freeze dataset, generated by Williams *et al.* [32] and Pop *et al.* [19]. Because of lower sequencing depth compared to Weinberg *et al.’s* data, the number of genes passing our selection criteria decreased to 625 genes for Williams and 212 for Pop (see **Materials and Methods**). We obtained an excellent correlation between our MER estimates for the two datasets (*r* = 0.92, p-value < 10^−5^, see Figure S11A. We also obtained a positive but less good correlation with Pop *et al.’s* dataset (*r* = 0.58, p-value < 10^−5^, see Figure S11B). As for the initiation rates, we also explained this less good correlation by the decrease in sequencing depth (which creates more sites with no footprints). By selecting the 30 codons with largest sample size used to compute the associated averaged codon elongation rate, the *r* coefficient indeed increased to 0.68.

Finally, since the differences in MER at different sites could be associated with tRNA availability variations [24], we further compared the MER and the codon tAI value [14,36], which reflects the codon usage bias towards the more abundant tRNAs in the organism, and found a positive correlation (*r* = 0.49, p-value < 10^−4^, see Figure 3C). Altogether, these results suggested that our estimates of the local elongation rates reflect tRNA-dependent regulation of elongation speed and that our estimates are consistent across different ribosome profile datasets.

### The impact of ribosomal interference on translation dynamics

The differences in the amount of ribosome interference between different genes could lead to significant biases when using the TE as a proxy for protein production rate. Using our results, we could quantify the production rate precisely, and thus relate it to the detected or total ribosome density. Simulating under our model and inferred parameters, we estimated the protein production rate using the particle flux: for each gene, we defined it as the rate at which a single ribosome reaches the end of the ORF and unbinds, leading to protein production. We examined the distribution of protein production rates (Figure 4A) and observed a range between 0.042 *s*^−1^ (5th percentile) and 0.12 *s*^−1^ (95th percentile), with median and standard deviation equal to 0.075 *s*^−1^ and 0.025 *s*^−1^, respectively. The protein production rate of a gene was generally lower than the corresponding translation initiation rate, due to an additional waiting time (~ 3 *s* on average) caused by ribosomal interference. Comparing the protein production rate with the detected-ribosome density (Figure 4A) gave a high correlation (Pearson’s *r* = 0.91, p-value < 10^−5^). However, we observed a super-linear increase of the production rate as the detected-ribosome density increased. Since our simulated detected-densities match the experimental TE measurements up to a normalization constant, this suggests that, because closely-stacked ribosomes are not included, the standard TE measure tends to underestimate the true protein production rate for large-TE genes. Using the total density of ribosomes (Figure 4A) instead of the detected-ribosome density improved the correlation (*r* = 0. 94, p-value < 10^−5^), but also led to a slight sub-linear trend, due to some saturation appearing when the initiation rate gets so high that elongation rates become limiting factors of translation.

**Figure 4.**
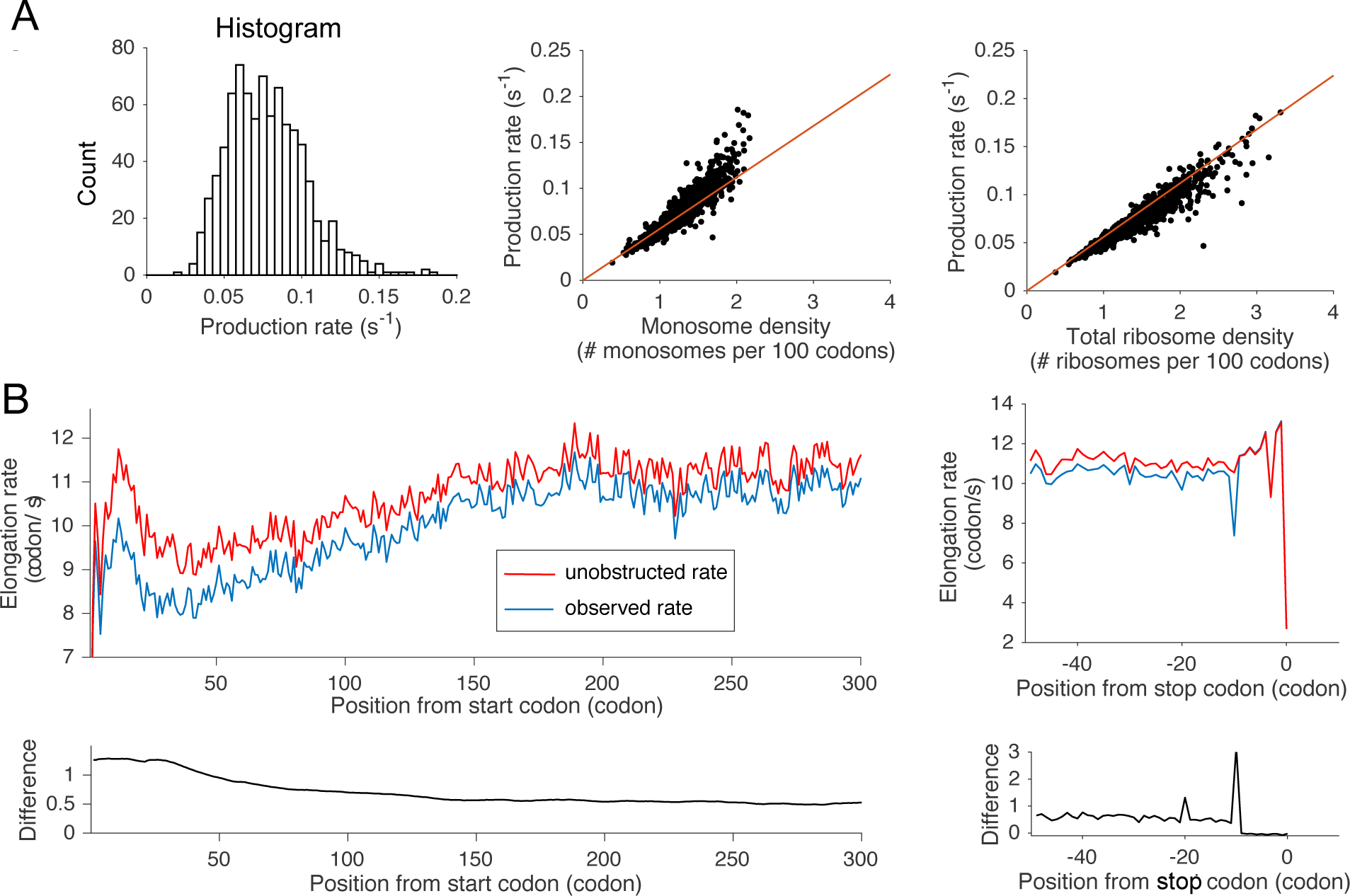
The impact of ribosomal interference on translation dynamics. **A**. Analysis of protein production. (Left) A histogram of protein production rates. (Middle) Comparison between the protein production rate and the detected-ribosome density obtained from simulations. In red, we plotted the simulated production rate as a function of ribosome density. The red line corresponds to the production rate when we assume no interference and a constant elongation speed of 5.6 codons/*s*, which was measured experimentally [7]. (Right) Comparison between the production rate and the total ribosome density density obtained from simulations. **B**. (Left) Position-specific elongation rates averaged over all transcript sequences, aligned with respect to the start codon. Plotted are the inferred unobstructed rate (in red) and the observed rate (in blue). The bottom plot shows the difference between the two curves. (Right) Similar plots as the ones on the left, when the transcript sequences are aligned with respect to the stop codon position.

To study how ribosomal interference affects the local ribosome dynamics, we examined the difference between the inferred elongation rates of our mathematical model (we call them *unobstructed* rates) and the effective rates given by the inverse of the average time spent at a particular position (we call them *observed* rates). Upon aligning all transcripts with respect to the start codon and averaging across the transcripts, we compared the average unobstructed rate at each position with the corresponding average observed rate (Figure 4B). Both curves showed an initial decrease to a trough located at codon position around 40, followed by a slow increase to a plateau. These variations were vertical reflections of the 5′ ramp obtained for ribosomal normalized density (Figure S12). Both unobstructed and observed rates initially increased from a very low rate (~ 3 codons/*s*) to a peak of 11.5 and 10 codons/*s*, respectively, located at position 10. They then decreased to a local minimum of 9 and 7.9 codons/*s*, respectively, before increasing again to a plateau around 11.5 and 10.9 codons/*s*, respectively. Furthermore, the gap between the unobstructed and observed rates generally decreased (Figure 4B, bottom plot) from 1.6 to 0.4 codons/*s* along the transcript, suggesting a decreasing impact of ribosomal interference on the translation dynamics. The reduction in the observed speed from the unobstructed elongation rate ranged from 5% (at the plateau) to 15% (between codon positions 10 and 20).

Aligning the transcript sequences with respect to the stop codon position and applying the same procedure, we observed a significant difference between the unobstructed and observed rates at codon position −10. The gap size is 3 codons/*s*, which amounts to 30% reduction from the unobstructed speed, while nearby sites have a regular level of 0.4 codons/*s*. This enhanced gap is likely induced by stalling at the stop codon. A smaller bump (1.3 codons/*s*) was also observed at codon position −20, reflecting the formation of a queue of three ribosomes.

### Variation of codon-specific mean elongation rates along the transcript

After studying the local dynamics of translation and quantifying the increase of elongation rates corresponding to the 5′ ramp of decreasing ribosome density, we investigated the possible determinants of such variation. The 5′ ramp of ribosome density has previously been attributed to slower elongation due to more frequent use of codons with low-abundance cognate tRNAs near the 5′-end [14]. However, this explanation has been argued to be insufficient [16,30], suggesting other mechanisms to cause the ramp.

To study whether the preferential use of slow codons can explain the variation of elongation rates along the transcript, we analyzed the positional distribution of different codons. To do so, we first grouped the codons (except stop codons) into five groups according to their mean elongation rates, and then plotted (Figure 5A) their frequency of appearance at each position in the set of genes we considered. At almost all positions, we found that the higher the mean elongation rate of a group, the higher the frequency of its appearance (the average frequency of appearance per codon type was 0. 25%, 0.9%, 1.6%, 1.9% and 2.25% for the five groups in increasing order of the mean elongation rate).

**Figure 5.**
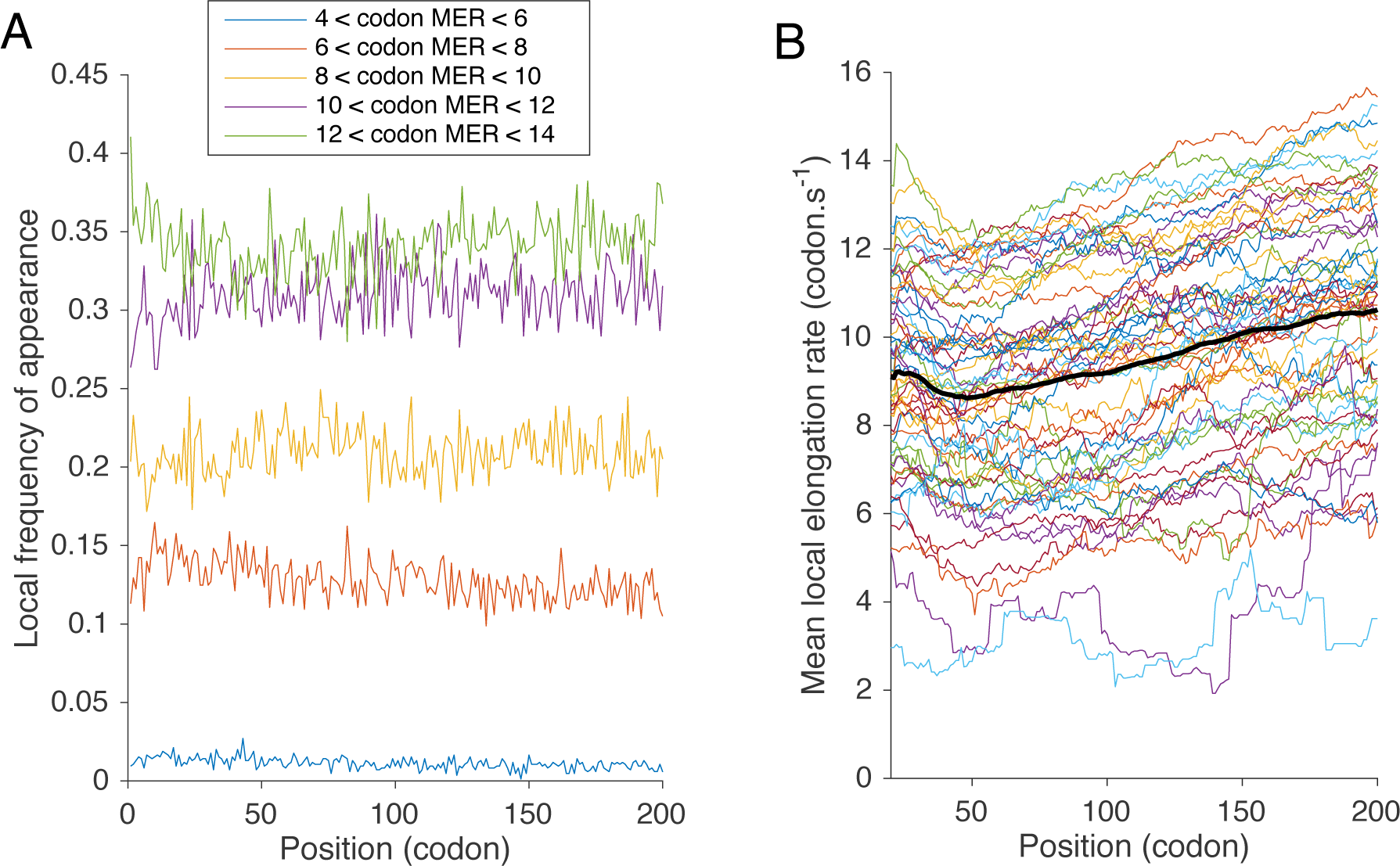
Heterogeneity of codon distributions and elongation speed along the transcript. **A**. Codon frequency metagene analysis. We grouped the codons (except stop codons) into five groups according to their mean elongation rates (MER) and plotted their frequency of appearance at each position in the set of genes we considered. The first group contained 4 codons with MER between 4 and 6 codons/*s*; the second group 13 codons with MER between 6 and 8; the third group 13 codons with MER between 8 and 10; the fourth group 16 codons with MER between 10 and 12; and the fifth group 15 codons with MER > 12. **B**. Smoothed mean elongation speed along the ORF for each codon type (stop codons are excluded). At each position *i*, we computed an average of codon-specific MER between positions *i —* 20 and *i* + 20. In black, we plot an average of the 61 curves.

Looking more closely at how these frequencies changed along the transcript between positions 50 and 200 (Figure S13A), we observed an increase in frequency for the fastest codons, while the opposite was true for slow codons. However, when we examined the associated positional variation in elongation speed by setting the elongation rate of each codon type at all positions to its corresponding average speed, we obtained an increase of 0.3 codons/*s* (Figure S14A). This increase was not large enough to explain the total variation observed at the 5′-ramp (approximately 2 codons/*s*). This result thus suggested the existence of other major factors influencing the elongation speed along the first 200 codons.

To confirm this hypothesis, we plotted the variation of average elongation speed for each codon type along the transcript sequence (Figure 5B), which displayed a range between approximately 2 and 14 codons/*s*. Also, for each position, we computed the mean deviation of each codon’s elongation rate from the codon-specific mean elongation rate. Figure S13B shows the results, which groups the codons according to their mean elongation rates, as done above.

Interestingly, we observed a general increase of the position-specific mean elongation rate from position 40 to 200 (corresponding to the ramp region). Weighting these variations by position-specific codon frequencies (Figure S14B), we found that the mean elongation rate from position 40 to 200 increases from approximately 9.5 to 11.5 codons/*s*, which gives an increase of 2 codons/*s*, comparable to what we previously observed in Figure 4B. We thus concluded that the major determinant of the 5′ translational ramp was not the codon distribution, but an overall increase of translational speed along the ORF.

### The major role of hydropathy and charge distributions of nascent polypeptides in explaining the positional variation of mean elongation rates

The above analyses suggested the existence of additional determinants that modulate local elongation rates and may explain the observed pattern of elongation rates along the transcript. We sought out to find these determinants using a statistical method.

Using molecular biology techniques, it has been demonstrated previously that electrostatic interactions between nascent polypeptides and the ribosomal exit tunnel can modulate elongation rates [37]. Motivated by this observation, we employed statistical linear models to identify specific features of the nascent polypeptide that affect elongation rates and to quantify the extent of their influence (see **Material and Methods**). We first analyzed Weinberg *et al*.’ s [16] data discussed above (850 genes). The dependent variable in each linear model was the position-specific mean deviation of elongation rates from codon-type-specific average elongation rates (the latter was obtained by averaging over all transcripts and positions). We used various features in our linear models, described in detail in **Material and Methods** and Figure S15.

About 40 or so amino acid residues can be accommodated within the ribosome [29], so we first considered codon positions 6 to 44 from the start codon in order to focus on the dynamics as the N-terminus of the nascent polypeptide chain makes its pass through the peptidyl transferase center (PTC) and the ribosomal exit tunnel. By optimizing the fit of linear models, we found that the PARS score in the window [9 : 19] downstream of the A-site is a statistically significant explanatory feature that is negatively correlated with the position-specific mean elongation rate in this region. This result is consistent with previous findings [30, 38] that mRNA secondary structure inhibits elongation near the 5′-end. This feature was generally more important for longer transcripts. We also found important regulatory features of the nascent polypeptide segment within the PTC and near the beginning of the exit tunnel. Specifically, when we used as additional features the mean number of charged amino acid residues and scanned linear models with different feature windows to obtain the best fit, we found that the number of positively charged residues in the window [1 : 11] and the number of negatively charged residues in the window [6 : 14] upstream of the A-site are important features with opposite effects; the former facilitates elongation, while the latter slows down elongation. These two charge features together with the PARS score explain 91% of the positional variation (Figure 6A) in the mean deviation of elongation rates in this region.

**Figure 6.**
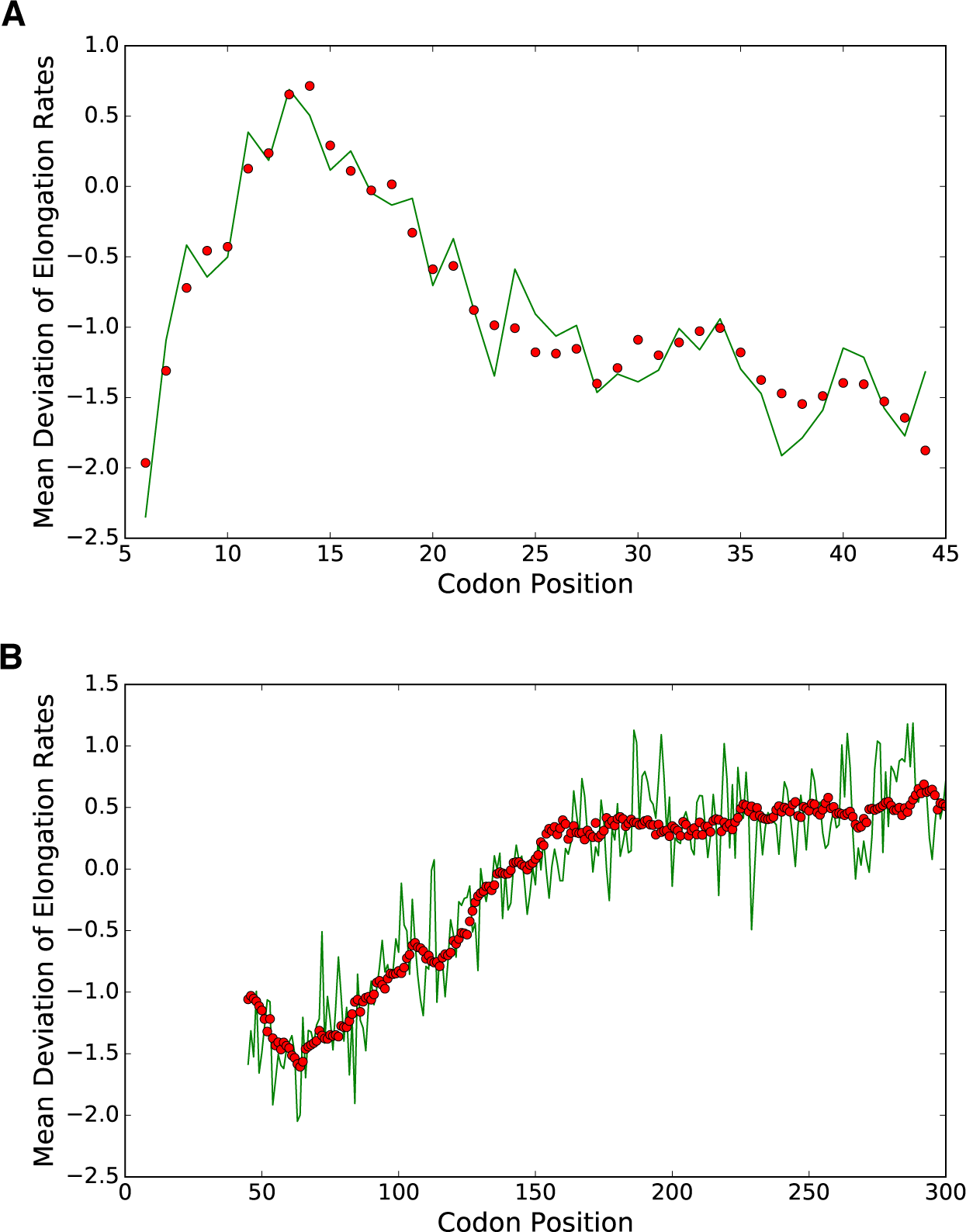
Linear model fits of the mean deviation of elongation rates for the data from Weinberg *et al.* [16]. The dependent variable is the mean deviation of elongation rates from codon-type-specific average elongation rates. Green lines correspond to the estimates from ribosome profiling data, while red dots correspond to our model fits based on a small (1 or 3) number of features. A. A fit for codon positions [6 : 44] obtained using three features: the mean PARS score in the window [9 : 19] downstream of the A-site, the mean number of negatively charged nascent amino acid residues in the window [6 : 14] upstream of the A-site, and the mean number of positively charged residues in the window [1 : 11] upstream of the A-site. The first two features had negative regression coefficients, while the last one had a positive regression coefficient. The coefficient of determination *R^2^* was 0.91 for this fit. B. A fit (*R*^2^ = 0.84) for the region [45 : 300] obtained using only a single feature: the mean hydropathy of the nascent peptide segment in the window [1 : 42] upstream of the A-site.

**W**e then tried to construct a linear model for codon positions 45 to 300. **W**e could not obtain a good fit only using explanatory features based on the PARS score and the number of charged residues. Surprisingly, we found that the hydropathy of the nascent polypeptide chain in the window [1 : 42] upstream of the A-site can alone explain 84% of the positional variation in the mean deviation of elongation rates in this region. This window [1 : 42] was determined by optimizing the fit of a linear model with hydropathy as the sole feature; the resulting fit is shown Figure 6B. This result implies that the more hydrophobic the nascent polypeptide segment is, the higher the mean elongation rate.

**W**e then took the above-mentioned features that we learned from analyzing the data from Weinberg *et al.* [16] and used them to fit the previously-mentioned ribosome profiling data for 625 genes from Williams *et al.* [32]. This led to fits with goodness comparable to the ones mentioned above: *R*^2^ = 0.86 for codon positions 6 to 44, *R*^2^ = 0.74 for positions 45 to 300, and *R*^2^ = 0.74 for the entire region between positions 6 and 300. A few factors potentially contributed to slightly lower coefficients of determination for Williams *et al.’s* data. First, 167 out of 625 genes in the dataset were shorter than 300 codons, while we excluded such genes when we analyzed Weinberg *et al.’s* data to eliminate the effects of ribosomal pausing near stop codons. Second, there are no RNA-seq data associated with the ribosome profiling from Williams *et al.*, so we could not refine the “naive” estimates of elongation rates for this dataset (see **Materials and Methods**).

### The charge features that modulate elongation rates are consistent with the electrostatic properties of the ribosome exit tunnel

To explain why the aforementioned windows of charge features got selected by our linear model of elongation rate variation, we studied the properties of the ribosome exit tunnel. To this end, we first extracted the ribosome tunnel coordinates and composition from cryo-EM data [39] using a tunnel detection algorithm [40] (see **Materials and Methods**, Figure 7 and Figure S16). The tunnel spans more than 80 Å and is composed of three regions: the upper region connected to the PTC, the constriction region (where two ribosomal proteins L4 and L22 reduce the width of the tunnel [29]), and the lower region connected to the exit (Figure 7A). Since our statistical analysis suggested that the presence of positive and negative charges in the upper region may respectively facilitate and inhibit elongation as the nascent polypeptide makes its initial pass through the tunnel (i.e., when the ribosome is translating the first ~ 40 codons), we aimed to study the longitudinal direction of the force that a charged particle would experience along the tunnel. Assuming the centerline of the tunnel to be approximately straight (fitting the 3D coordinates of the tunnel with a straight line gives a R-squared value of 0.985, see Figure S17), we can actually reduce the dynamics of a particle inside the tunnel into a one-dimensional diffusion process, driven by two potentials (see **Materials and Methods**): one associated with the entropy of the system related to the cross-sectional area, and the other with the electrostatic potential along the tunnel, created by the charged constituent RNAs and proteins of the ribosome. The local signs of the gradients of these potentials determine whether the movement of a charged particle towards the exit of the tunnel is facilitated or inhibited.

**Figure 7.**
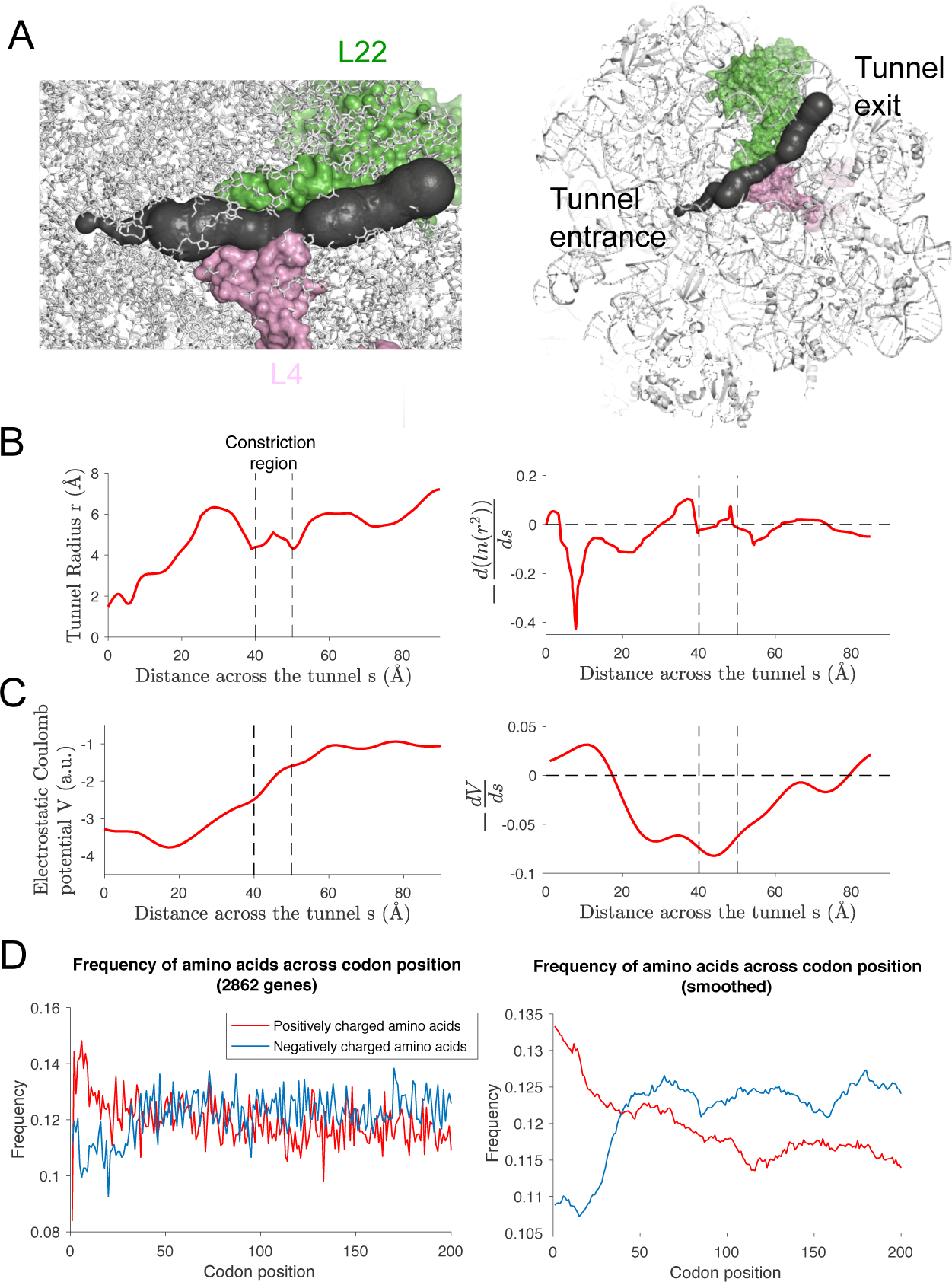
Biophysical properties of the ribosome exit tunnel. **A**. The figure on the left illustrates the exit tunnel (in black) and ribosomal proteins L4 (in pink) and L22 (in green) surrounding the constriction region. We extracted the tunnel geometry from the cryo-EM structure of the ribosome large subunit [39] illustrated on the right (side view, see also Figure S16). **B.** (Left) The variation of the tunnel radius *r* along the tunnel. (Right) The negative gradient of ln(r^2^) (smoothed over a 10 Å window). Note a region of large negative “entropic” potential [73] in the upper region of the exit tunnel. **C**. (Left) The variation of the electrostatic Coulomb potential induced by ribosomal RNA and proteins within a radius of 20 Å from the center of the tunnel. (Right) The negative gradient of the potential (smoothed over a 10 Å window), notably showing a region of positive electric field (pointing towards the exit) in the region 0 ~ 18 Å. D. (Left) Position-specific average frequencies of positively (red) and negatively (blue) charged amino acids averaged over all genes of length ≥ 200 codons. (Right) The frequency curves smoothed by averaging over a 10-codon window. Compared to other parts of the transcript, the frequency of positive (negative) amino acids is significantly higher (lower) in the first ~25 codons.

By studying the entropic force (Figure 7B), we found that the gradient was small in the constriction and lower regions but positive and much larger in the first half of the upper region (first ~ 20Å). Thus, a particle crossing this region will experience a strong entropy barrier due to the increase of radius in the tunnel. For the electrostatic force, due to the heterogeneity of the solvent inside the tunnel [41], it is difficult to accurately compute the electrostatic potential inside the tunnel using classical numerical methods [42]. Therefore, we simply approximated the electrostatic potential by the Coulomb potential generated by the ribosome, obtained at any point of the centerline by summing the charges in the ribosomes (coming from negatively charged phosphate groups and charged amino acids) weighted by the inverse of the distance to the point. Using this approximation, we found that the electric field is outward-pointing in the first half of the upper region (0-20 Å) (Figure 7C). As the tunnel can accommodate approximately 40 amino acids [29], we concluded that this is consistent with our results that the presence of positively charged residues in the first 11 amino acids of the nascent polypeptide tends to facilitate elongation, whereas negatively charged residues between positions 6 and 14 tend to slow it down (see Discussion below). Moreover, these results suggest that the presence of positively charged amino acids should help in the first steps of elongation to move the N-terminus of the nascent polypeptide across the entropy barrier and towards the tunnel exit.

Finally, we studied whether there is an evolutionary signature that is consistent with our finding regarding the specific role that electrostatic interaction plays in modulating elongation speed as the N-terminus of the nascent polypeptide makes its way through the tunnel. To this end, we looked at the position-specific frequency of positive and negative amino acid charges over an extended set of genes (all 2862 genes of length ≥ 200 codons) in the first 200 codons of the ORF. As shown in Figure 7D, the frequency of positive charges was much larger in the first 40 codons, followed by a gradual decrease to a plateau (~ 12% at position 50 and ~ 11% at 200). In contrast, we observed a stark depletion in the frequency of negative charges in the beginning of the ORF, followed by an increase to a plateau starting around position 40 (~ 12.5%); Tuller *et al.* [30] also made a similar observation. Overall, these patterns are consistent with the results of our statistical analysis and our hypothesis that evolution has tried to optimize charge distributions in the beginning of the nascent polypeptide to facilitate elongation as the N-terminus make its way through the exit tunnel.

## DISCUSSION

### Difference of our method from previous methods

We used probabilistic modeling of the translation dynamics to dissect the different determinants of elongation speed, and developed an efficient, simulation-based inference algorithm to estimate transcript-specific initiation and local elongation rates from ribosome profiling data. The first step of our inference procedure is similar to the method introduced by Ciandrini *et al.* [34], which uses a TASEP-based model to infer gene-specific initiation rates; in our case we use transcript- and position-specific elongation rates, by taking the inverse of the profile density, whereas Ciandrini *et al.* use codon-specific elongation rates derived from tAI values, common to all transcripts and positions. As Ciandrini *et al.’s* method uses only the ribosome density from polysome profile, it assumes that the elongation rate depends only on the codon identity at the A-site, neglecting other determinants that we notably found in our study to explain the elongation rate variability. While our estimates of initiation rates were of similar order and positively correlated, it is interesting to note that the correlation decreases for genes with higher initiation rates. This could be due to the additional information (specifically, the positions of ribosomes on mRNAs) provided by ribosome profiling that are missing in their method. For example, Ciandrini *et al.* assumed that termination is a fast process, although our results show the contrary. This suggests that they may underestimate the impact of interference, causing them to overestimate the initiation rate necessary to reach a given density. This is consistent with the fact that their estimated initiation rates are in general slightly larger than our estimates.

Gritsenko *et al.* [20] proposed another TASEP-based approach to estimate initiation and elongation rates from ribosome profiling data. However, in their approach only 61 parameters of elongation are estimated by minimizing an objective function over all the profiles. The observed difference between tAI-based and their fitted elongation rates led them to conclude that additional unknown factors, possibly arising from larger sequence context, are shaping elongation rates. Our results illustrate this point more precisely and show that some specific properties of the nascent polypeptide can explain the variability of elongation rates that we observed across different transcripts and codon positions.

As detailed in a recent review by Zur and Tuller [43], there has been many previous studies using computational tools to infer and predict the dynamics of mRNA translation using biophysical modeling [10, 14, 15, 19, 20, 30, 43], but with contradictory results. To our knowledge, the models proposed to date, except for the one considered by Tuller *et al.* [30], have been developed under the assumption that elongation rates are not influenced by the sequence context surrounding the A-site, while our results suggest that elongation rates are modulated by the nascent polypeptide interaction with the exit tunnel that depend on the context of ~ 40 codons preceding the A-site. Interestingly, the model proposed by Tuller *et al.* also takes into account amino acid charges and mRNA folding energy, but their conclusion regarding the impact of the charges on the elongation rate strongly differs from what our results suggest (see below). Furthermore, their model does not include hydropathy, which we show plays a major role in regulating elongation speed once the N-terminus has escaped the ribosome exit tunnel.

### Potential technical artifacts

When combining a biophysical modeling approach with ribosome profiling data, another source of complication comes from technical artifacts in the data. In particular, cycloheximide pre-treatment, used to immobilize ribosomes, can lead to substantial codon-specific biases [16, 44, 45]. The use of flash-freeze technique alleviates some of these problems, and allows one to obtain ribosome-footprint profiles and mRNA abundances that more faithfully reflect the translation dynamics [16]. Andreev *et al.* [12] recently described several important artifacts and biases associated with ribosome profiling data that affect the representation of translation dynamics. The experimental protocol used for the main flash-freeze data [16] considered in this paper minimizes some of the biases (such as sequence biases introduced during ribosome footprint library preparation and conversion to cDNA for subsequent sequencing, and mRNA-abundance measurement biases and other artifacts caused by poly(A) selection). In addition, our method allowed us to correct other important biases related to TE measurements and depletion of stacked ribosomes from selecting only ~ 30 nt fragments.

### Undetected ribosomes and the extent of interference

Our quantification of interference was made upon selecting a model of ribosome profiling with no detection of closely-stacked ribosomes. This model was selected over other models with partial or total detection of closely-stacked ribosomes, as the simulated total densities under this model produced the best agreement with polysome profiling data from Arava *et al.* [33]. We note that comparing TEs with another polysome profiling dataset [35] yields a smaller scaling constant (0.7 instead of 0.82), which would lead to lower initiation rates and less interference. However, for this dataset, we still observe a decrease of the fitting constant for genes with higher density (0.58). In another study [46], we investigated different gap sizes using a refined analysis of the TASEP to study the relation between the total density and the density of isolated ribosomes for a specific isolation distance. The best match was obtained for a distance of 2 or 3 codons, in agreement with the model used here. Interestingly, applying our inference procedure also shows that our estimated elongation rates are in vast majority the same as the naive estimates obtained from inverting the profile density (see **Materials and Methods**). Hence, using an alternate model accounting for possible detection of closely-stacked ribosomes would not modify the main conclusions of our work, notably in regards to the major determinants of the elongation rate. It is also possible that for other datasets and other organisms, differences in experimental protocols and nuclease digestion conditions [47] could affect the fraction of detected stacked-ribosomes. Although it has been shown that ribosome profiling in yeast yields reproducible datasets with minimal variations [48], analyzing datasets from other organisms could possibly require using an alternate model.

### The major determinants of the elongation rate

Our inferred rates from flash-freeze data showed that the elongation rate is indeed modulated by the decoded codon located at the A-site of the ribosome and the corresponding tRNA availability. The positive correlation between the codon-specific mean elongation rate and the translation adaptation index, which has been used as a proxy for codon-specific decoding rate [10, 34, 36, 43], supports the hypothesis that tRNA abundance and codon usage co-evolved to optimize translation rates [14, 49].

However, our refined analysis of the distribution of codon-specific elongation rates showed that tRNA availability is not sufficient to fully explain the observed translational speed variation. In particular, the 5′ translational ramp variation cannot be sufficiently explained by the change of frequencies of slow and fast codons across the transcript sequence, contrary to what was previously suggested [14, 30]. Indeed, subsequent studies showed that specific configurations of amino acids along the nascent polypeptide segment within the exit tunnel can contribute to a slowdown or arrest of translation [17, 18, 27, 29, 50, 51]. An earlier study [37] proposed that electrostatic interactions of nascent polypeptides with the charged walls of the ribosomal exit tunnel could be one of the possible mechanisms of modulation of elongation speed, suggesting that positively charged amino acids slow down translation [17, 30, 37]. We note, however, that these studies did not focus on the initial stage of elongation. In contrast, our results suggest that while the N-terminus of the nascent polypeptide has not exited from the tunnel, positively charged amino acids in specific parts of the polypeptide actually facilitate the elongation speed, while the opposite is true for negatively charged amino acids. Once the N-terminus has exited the tunnel, the hydropathy of the part of the nascent polypeptide within the ribosome plays a major role in governing the elongation rate variation. These features were selected by statistically optimizing the fit of the linear model to position-specific mean elongation rates in a large region, which included the 5′ ramp.

**W**e note that Tuller *et al.* [30] also employed linear regression to fit the 5′ ramp and developed a model that includes various features such as the tAI value, the total charge of the amino acid residues coded by the 13 codons upstream of the A-site, and the 5′ mRNA folding energy downstream of the A-site. Although these features seem similar to ours, the results are quite different. Indeed, while Tuller *et al.* suggested that the ribosome density along the first 300 codons can be explained by these features, our statistical analysis showed that the codon distribution (associated with tAI variations) does not significantly contribute to the ramp variation (Figure S14A). Further, we obtained that the amino acid charges and mRNA secondary structure only explain the elongation rate variation during the first stage of elongation (~ first 45 codons, Figure 6A), while the remaining part of the ramp is mainly explained by the hydropathy score of the nascent polypeptide located within the exit tunnel (Figure 6B). These results are actually consistent with Tuller *et al.’s* finding, as they showed a poor correlation between their model and the ribosome density after the first 50 codons [30]. Moreover, as previously said, the detailed impact of the charges on the elongation dynamics is the opposite of what we found, as they suggested that the ribosome density is positively correlated with the charges (in other words, positive charges are negatively correlated with the elongation rate, and thus slow down elongation).

There are several possible explanations for the differences between Tuller *et al.* ’ s results and ours. First, since their analyzed footprints came from an experiment that used cycloheximide pre-treatment, their fit for the first 50 codons could be affected by data artifacts associated with the chemical treatment. In addition, the features used in their model were hand picked, while ours were obtained through an optimization procedure. Finally, they fitted a smoothed version of the normalized average ribosome footprint density along the transcript (see Figure S12A), whereas we fitted the position-specific deviation from the mean elongation rate. Smoothing could be undesirable, as we showed that some part of the density variation in this region is due to interference (Figure 4).

### Possible biophysical explanations

There are reasonable biophysical explanations for the particular set of features selected by our statistical analyses. In order for the ribosome to translocate from one site to the next, the nascent polypeptide has to be displaced to liberate enough space for the chain to incorporate the next amino acid. The associated force needed to achieve this process is constrained by the biophysics of the tunnel, which is known to be charged, aqueous, and narrow [29, 37, 52]. When the nascent polypeptide has not yet exited the tunnel (i.e., the first 40 residues), our statistical analysis found that charged amino acid residues near the A-site play an important role in governing the elongation dynamics. Furthermore, we found that positive charges and negative charges have opposite effects: the former facilitates the elongation speed, while the latter inhibit it. This finding is consistent with the electrostatic properties of the tunnel. Specifically, our estimations of the net local charge across the tunnel suggests that the electric field induced by the potential points outward (i.e., away from the PTC) near the beginning of the tunnel (Figure 7C), in agreement with what our linear model predicts. Previous measurements of the electrostatic potential inside the tunnel [53] were also consistent with our findings, suggesting a decrease of the potential from the PTC along the upper region. Hence, positively charged residues near the beginning of the tunnel will experience an electrostatic force pointing outward, thereby facilitating the movement of the polypeptide chain through the tunnel. Moreover, we showed that in the context of studying particle diffusion inside the tunnel, the increase of radius across the tunnel in the upper region (Figure 7B) creates a strong entropic barrier. This barrier can be compensated by the electrostatic potential if the particle is positively charged, explaining the specific selection of positively charged amino acids in the upper region when the nascent polypeptide makes its initial pass through the tunnel. The opposite applies to negatively charged residues, with an effect of inhibiting the movement of the chain.

Averaging the charged amino acid frequency over all transcripts of length ≥ 200 codons, we found that there is a starkly elevated amount of positively charged amino acids in the first 25 codons, while the opposite is true for negatively charged amino acids (see Figure 7D). These patterns are consistent with our proposed role of positive and negative charges, and suggest that evolution has tried to optimize charge distributions to facilitate the translation dynamics as the nascent polypeptide makes its initial pass through the exit tunnel.

Another important feature, which to our knowledge has not been previously noted as a major determinant of elongation speed, is the hydropathy of the polypeptide segment within the PTC and the exit tunnel. A possible explanation for the impact of hydropathy on the elongation rate is that since the tunnel is aqueous [29] and wide enough to allow the formation of α-helical structure [52], the hydrophobicity (which is an important factor driving compactness and rigidity [54]) of the polypeptide segment inside the ribosome consequently drives the amount of force needed to push the chain up the tunnel. This result may seemingly be in contradiction with the fact that membrane proteins, known to be more hydrophobic, are generally more difficult to express. Analyzing a subset of 89 membrane protein genes [55] in our main dataset (see Figure S19), we indeed found a significant decrease in TE, but this decrease is actually explained in our model by lower initiation rates. Interestingly, while we observed a larger average hydropathy score for membrane proteins, we also found significant variation in hydropathy along the sequence, with a larger increase from position 40 to 200 compared to the increase observed in other proteins (Figure S18). Interestingly, the 5′ ramp pattern is also present in the 89 membrane protein-coding genes (Figure S19). The larger amplitude of the ramp also suggests a larger increase in the elongation rate, in agreement with our original finding regarding the impact of hydropathy on the elongation rate.

Shalgi *et al.* [56] recently showed that translation elongation pausing around position 65 is associated with hydrophobic N-termini, under stress conditions reducing interactions between the ribosome and the Hsp70 family of chaperones. Our analysis suggests that, more generally, the variation of the hydrophobicity of the nascent polypeptide inside the exit tunnel (Figure S18) could explain the variation of the elongation rate over a large region following codon position ~ 50, and notably the trough observed around the same position as in Shalgi *et al.* While variation in translation rates could play a functional role in regulating co-translational folding of the nascent polypeptide chain [57], our results on the impact of hydropathy suggest that this link is more complex in that the folding (or pre-folding) in turn can actually alter the rate of translation. Interestingly, the observed variation in the mean hydropathy score along the transcript (Figure S18) suggests that the elongation speed is regulated at different stages of the polypeptide assembly and folding. The selection of different determinant features for different stages of translation also suggests that the movement of the polypeptide inside the tunnel is driven by two distinct biophysical mechanisms: First, when the polypeptide chain has not yet exited the tunnel, electrostatic interactions in the upper part of the tunnel play a major role in regulating the movement of the chain down the exit tunnel. Second, when the polypeptide has reached a certain length and its N-terminus has exited the tunnel, it is the structure of the chain itself (which we captured through the hydropathy) that determines its movement through the tunnel.

### Possible limitations

One of the possible limitations of our present approach is that it does not take into account possible translational regulatory events which could be specific to some genes (such as co-translational translocation of membrane proteins, which may slow down the translation rate [58]) or sequence motifs (such as arrest sequences inducing mRNA cleavage [59]). Another regulatory mechanism is ribosomal drop-off, which has been shown to occur for some sequences that lead to pausing or under specific stress conditions [22]. Under a non-stress environment, it has been hypothesized that there exists a “basal” drop-off rate (defined as the probability per site for a ribosome to drop-off from the transcript it is translating) and it has been estimated to be on the order of 10^-4^ per codon in *E. coli*, assuming sequence-specific features to be well averaged out. This has led to the hypothesis that ribosomal drop-off could explain the ramp variations [22]. Interestingly, using the same method as in Sin *et al.* [22] (see Section 7 of Supporting Information), we came to different estimates of the drop-off rate for the ramp region and the rest of the transcript (Figure S20A-B). More precisely, while the drop-off rate we estimated for the region outside the ramp was consistent with the drop-off rate estimate (3.7 × 10^−4^) from Sin *et al.*, the estimation procedure led to an unrealistically large drop-off rate in the ramp region (0.002, which leads to a survival probability of only 0.67 after 200 translated codons). Incorporating the basal drop-off rate of 3.7 × 10^−4^ per codon into our inference procedure did not significantly change our rate estimates (see Figure S20C). Another possible drawback of our work is that our main analysis was carried out on a subset of only 850 genes, which were selected to assure sufficiently high local coverage-depth of footprints. To confirm that our results and conclusions did not suffer from any potential biases due to such filtering, we analyzed the ramp pattern and the codon-specific elongation rates obtained from a larger dataset (2862 genes) consisting of all genes of length ≥ 200 codons. Comparison with our original results (see Figure S21) did not show any significant difference, suggesting that our overall conclusions are robust and applicable to a broader level.

Finally, we have tried to apply our method to individual mRNA sequences, but could not obtain significant results. We believe that aside from some logical issues related to noise at the single transcript level, the very simple features that we used are reflecting some modulation at a higher level, once all the interactions that affect the elongation rates are averaged over all the transcripts. For example, the simple hydropathy score we considered may be able to capture that the structure is important, but it cannot reflect the folding of the individual polypeptide at a specific location. Similarly, our approximation of the electrostatic potential is insufficient to quantify the amount of electrostatic force applied to the polypeptide, which should be a much more complex function of the spatial distribution of charges and amino acids [28]. It is also possible that some features (for example the presence of poly proline) [18] can specifically affect the elongation rate, but cannot be detected as a global feature obtained after averaging. Finally, using a linear model to combine these features may be too simplistic and insufficient to capture how a specific sequence context can locally affect the elongation rate. Overall, our results suggest that a better understanding of the biophysical interactions between the tunnel and the nascent chain is needed to study the translation dynamics at early stage.

### Conclusion

In summary, our results show how the time spent by the ribosome decoding and translocating at a particular codon site is governed by three major determinants: ribosome interference, tRNA abundance, and biophysical properties of the nascent polypeptide within the PTC and the ribosome exit tunnel. It is quite remarkable that using a linear model with only few features allowed us to fully and robustly capture the variations of the average elongation rate along the transcript sequence. The results from our statistical analysis suggest that the translation elongation dynamics while the nascent polypeptide is initially passing through the ribosome exit tunnel is rather different from the elongation dynamics after the N-terminus has escaped the tunnel, and that different biophysical mechanisms modulate the elongation speed in the two stages. In addition to these overall determinants, our study also demonstrated the importance of mRNA secondary structure in the first 40 codons and a pausing of the ribosome at or near the stop codon, suggesting that additional local mechanisms may play a role in modulating translation in specific parts of a transcript sequence.

Since the ribosome structure is highly conserved [60, 61], we believe that our results can be generalized to other organisms. However, applying the present method to other datasets to estimate translation rates may not be straightforward. Differences in experimental protocols and nuclease digestion conditions could affect the data substantially [47], and in particular make the fraction of detected stacked-ribosomes to vary across different datasets and organisms. However, combining polysome profiling, TE measurements, and different profile simulation models (with different probabilities of detecting stacked-ribosomes) can allow one to estimate the proportion of undetected stacked-ribosomes (as done in Figure 2 and Figure S8), and also enable the estimation of translation rates upon selecting the best model. Finally, a natural extension of our work is to investigate in more detail, based on the above findings, the determinants of translation at the individual transcript level. To do so, a more detailed analysis and modeling of the nascent polypeptide within and immediately outside the exit tunnel is needed, to reveal how a specific amino acid sequence can affect the translation rate through possible interactions or co-translational folding [57, 62, 63].

## MATERIALS AND METHODS

### Experimental dataset

We used publicly available data in our analysis. The flash-freeze ribosome profiling data from Weinberg *et al.* [16] can be accessed from the Gene Expression Omnibus (GEO) database with the accession number GSE75897. The accession number for the flash-freeze data from Williams *et al.* [32] is GSM1495503 and the one from Pop *et al.* [19] are GSM1557442 (RNA-seq) and GSM1557447 (ribo-seq). The method used to map ribosome footprint reads is described in Weinberg *et al.* [16] (or further details on the choice of these datasets, see Section 5 of **Supporting Information**). To be able to determine normalization constants (detailed below) without being biased by the heterogeneity of translational speed along the 5′ ramp and to obtain robust estimates of the steady-state distribution, we selected among the pool of 5887 genes the ones longer than 200 codons and for which the average ribosome density was greater than 10 per site. For the Weinberg *et al.* dataset this led to a set of 894 genes, to which we applied the first step of our inference procedure (described below) to produce an estimate of the initiation rate. The algorithm converged for 850 genes, and the main results presented in this paper are based on those genes. For the Williams *et al.* dataset, the same procedure gave 625 genes. For Pop *et al.* it gave 212 genes.

### Mapping of the A-site from raw ribosome profile data

To map the A-sites from the raw short-read data, we used the following procedure: We selected the reads of lengths 28, 29 and 30 nt, and, for each read, we looked at its first nucleotide and determined how shifted (0, +1, or -1) it was from the closest codon’s first nucleotide. For the reads of length 28, we assigned the A-site to the codon located at position 15 for shift equal to +1, at position 16 for shift equal to 0, and removed the ones with shift —1 from our dataset, since there is ambiguity as to which codon to select. For the reads of length 29, we assigned the A-site to the codon located at position 16 for shift equal to +0, and removed the rest. For the reads of length 30, we assigned the A-site to the codon located at position 16 for shift equal to 0, at position 17 for shift equal to —1, and removed the reads with shift +1. Such assignments of the A-site leaves about 80% of the total data.

### Estimation of detected-ribosome densities from translation efficiency measurements

Translation efficiency measurements were used to compute the average density of detected ribosomes. Since translation efficiency is given by the ratio of the RPKM measurement for ribosomal footprint to the RPKM measurement for mRNA, it is proportional to the average density of detected ribosomes.To estimate the associated constant for each gene of our dataset, we used the measurements of ribosome density from Arava *et al.* [33] (the dataset contains 739 genes, with 588 common with Weinberg *et al.* [16]). For genes with a ribosome density of less than 1 ribosome per 100 codons (110 genes in total), we fitted the translation efficiency as a function of the density to a linear function and divided all the TEs by the coefficient of this fit to obtain estimates of the detected-ribosome density.

### Estimation of 5′-cap folding energy

The 5′-cap folding energy associated with each gene of our dataset was taken from Weinberg *et al.* [16], who used sequences of length 70 nt from the 5′ end of the mRNA transcript and calculated the folding energies at 37°C using RNAfold algorithm from Vienna RNA package [64].

### Estimation of RNA secondary structure (PARS score)

To quantify RNA secondary structure at specific sites, we used the parallel analysis of RNA structure (PARS) scores from Kertesz *et al.* [65]. It is based on deep sequencing of RNA fragments, providing simultaneous *in vitro* profiling of the secondary structure of RNA species at single nucleotide resolution in *S. cerevisiae* (GEO accession number: GSE22393). We defined the PARS score of a codon by averaging the PARS scores of the nucleotides in that codon.

### Mathematical modeling of translation

To simulate ribosome profiles, we used a mathematical model based on the totally asymmetric simple exclusion process (TASEP) [31, 66]. Compared with the original TASEP, our model included additional features accounting for the heterogeneity of elongation rates and physical size of the ribosome. We assumed that each ribosome has a footprint size of 30 nucleotides (i.e., 10 codons) and that the A-site is located at nucleotide positions 16-18 (from the 5′ end) [67]. Protein production consists of three phases: First, a ribosome enters the ORF with its empty A-site at the second codon position; the waiting time follows an exponential distribution and we define its rate as the initiation rate. Subsequently, a ribosome is allowed to move forward one codon position if this movement is not obstructed by the presence of another ribosome located downstream. As the dynamics of a ribosome along an mRNA transcript can be seen as a Markov jump process, the associated conditional hopping time at each site is exponentially distributed, with its rate defined as the elongation rate at the site. When a ribosome eventually reaches a stop codon, it unbinds at an exponential rate (for simplicity, we also refer to this as an elongation rate), which eventually leads to protein production. By simulating under this model with given initiation and position-specific elongation rates, we can sample ribosome positions at different times and thereby approximate the marginal steady state distribution of ribosome positions (for further details, see Section 1 of Supporting Information). In practice, we sampled ~ 3 × 10^4^ ribosome positions for each gene studied.

### Definition of closely stacked ribosomes

During simulation we monitor the distance between consecutive ribosomes along the transcript. Since experimental “disome” fragments (i.e., footprints covering two stacked ribosomes) were shown [11] to protect a broad range of sizes below ~ 65 nt, in our simulations we defined closely stacked ribosomes to occur when the distance between the A-sites of consecutive ribosomes is ≤ 12 codons (i.e., free space between the ribosomes is ≤ 2 codons).

### Inference procedure

A detailed description of our inference procedure with examples is provided in Section 3 of Supporting Information. Briefly, for given experimental ribosome profile and detected density (average number of detected ribosomes occupying a single mRNA copy), our inference procedure for estimating transcript-specific initiation and local elongation rates of the assumed TASEP model consists of two steps (Figure 1 B). 1) First, we approximate the position-specific elongation rate by taking the inverse of the observed footprint number (such approximation is valid when there is no ribosomal interference, see Section 3 of Supporting Information), and then use simulation to search over the initiation rate that minimizes the difference between the experimental detected-ribosome density and the one obtained from simulation. 2) Then, simulating under these naive estimates, we compare the simulated ribosome profile with the experimental one and detect positions, called “error-sites”, where the absolute density difference is larger than a fixed threshold. If error-sites are detected, we first consider the one closest to the 5′-end. We jointly optimize the elongation rates in a neighborhood of this error-site and the initiation rate to minimize the error between the simulated and the observed profile. With these new parameters, we then re-detect possible error-sites located downstream and repeat the procedure until there are no more error-sites located downstream to correct.

Because the profile and average density are invariant to a global scaling of the initiation and elongation rates, the parameters obtained needed to be normalized to get the rates in appropriate units. We normalized the rates such that the global average speed measured by simulations between position 150 and the stop codon is 5.6 codons/*s*, as measured experimentally [7]. We restricted our analysis to genes longer than 200 codons so that this normalization procedure is not biased by the heterogeneity of translational speed along the 5′ ramp.

For given initiation and position-specific elongation rates along the transcript, obtaining an analytic formula for the protein synthesis flux is often difficult, if not impossible, due to potential interference between ribosomes occupying the same transcript [68–70]. However, since translation is generally limited by initiation, not by elongation, under realistic physiological conditions [15, 71], typically only a few sites were affected by interference (Figure S2A). This allowed us to cope with the high dimensionality of the model space and obtain estimates of rate parameters that produced excellent fit to the experimental data (Figure S1).

### Linear model of elongation rate variation

After inferring the elongation rates, we employed statistical linear models to fit the positional variation of the mean elongation rate. For a given position *i*, our dependent variable *y_i_* was given by

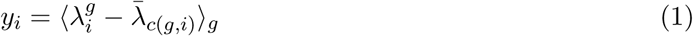

where 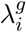 denotes the inferred rate for position *i* of gene *g, c*(*g*,*i*) the codon at position *i* of gene *g*, and 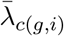 the codon-type-specific average elongation rate obtained by averaging over all transcripts and positions. The notation 〈·〉_*g*_ denotes taking an average over the gene set. **W**e considered linear models of the form

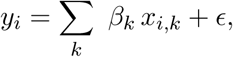

where *x_i,k_* are explanatory variables for position *i* obtained by averaging over all genes in our dataset, *β_k_* are regression coefficients, and *∊* is a noise term. The explanatory variables *x_i,k_* were obtained from considering some biophysical properties of the nascent polypeptide and the sequence context (see below). We tested various models with different choices of explanatory variables and tried to find the best fitting model.

One kind of explanatory variable we considered is the average PARS score [65], which reflects the existence of mRNA secondary structure, over a window [a : b] located downstream of the A-site: for a given A-site i, the PARS score window [a : b] refers to positions from *i* + *a* to *i* + *b*. The other features were related to the nascent polypeptide properties, namely total positive charges, negative charges, and hydropathy scores [72]. For these features, we used different windows located upstream of the A-site; in this case, a window [a : b] for a given A-site *i* refers to positions from *i — b* to *i — a.* The negative (positive) charge feature was obtained by computing the number of glutamic and aspartic acids (arginines and lysines) located upstream of the A-site in the specified window. An illustration of how the variables are computed is provided in Figure S15.

### Ribosome cryo-EM data and exit tunnel extraction

The ribosome crystallographic structure from Schmidt *et al.* [39] (Protein Data Bank ID 5GAK, resolution ~ 3.88 Å) was used to study the ribosomal exit tunnel, and the structure was visualized using Pymol. We extracted the tunnel coordinates using MOLE 2.0 software [40], and used custom python and Matlab scripts to compute the radius and charge properties.

### Diffusion of a particle in a three-dimensional structure with a varying crosssection size

The diffusion of a particle in a three-dimensional structure with a varying cross-section size can be treated as a one-dimensional process with an entropy barrier, described by the so-called Fick-Jacobs diffusion equation [73]. The Fick-Jacobs equation has the same structure as the Smoluchowski equation for diffusion in a one-dimensional potential [73], where the potential *S*(*x*) arises from the entropy along the tunnel, determined by the cross-sectional area as

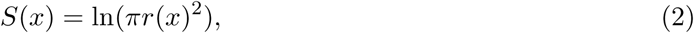

where *r*(*x*) is the radius of the cross-section at position *x* along the tunnel.

### Software implementation

Simulation of translation and our inference algorithm were implemented in Matlab. We simulated the model using the next reaction method [74] derived from the Gillespie algorithm, which at each step samples the next event (initiation, elongation, or termination) and the associated time based on the current ribosome occupancy (see Section 2 of **Supporting Information**). To simulate a ribosome profile of size *N*, we first simulated ~ 10^4^ steps for burn-in. Then, after a fixed interval of subsequent time steps, we randomly picked one occupied A-site (if there is one) and recorded it as a footprint location; this sampling scheme was iterated until we obtained *N* footprints. Protein production flux was obtained by computing the ratio between the number of ribosomes going through termination and the total time.

### Supporting Information

Description of the probabilistic model, simulation algorithm, inference procedure, and estimation of drop-off rates.

## ACKNOWLEDGMENTS

We thank Oana Carja, Joshua Plotkin, Premal Shah, and David Weinberg for useful discussions and for providing with the data analyzed in this paper. We also thank Barry Cooperman, Carol Deutsch, Steve Harvey, and Kim Sharp for helpful discussions on biophysics. This research is supported in part by National Science Foundation (NSF) CAREER Grant DBI-0846015, a Math+X Research Grant from the Simons Foundation, and a Packard Fellowship for Science and Engineering. YSS is a Chan Zuckerberg Biohub investigator. The funders had no role in study design, data collection and analysis, decision to publish, or preparation of the manuscript

## SUPPLEMENTARY FIGURES

**Figure S1.**
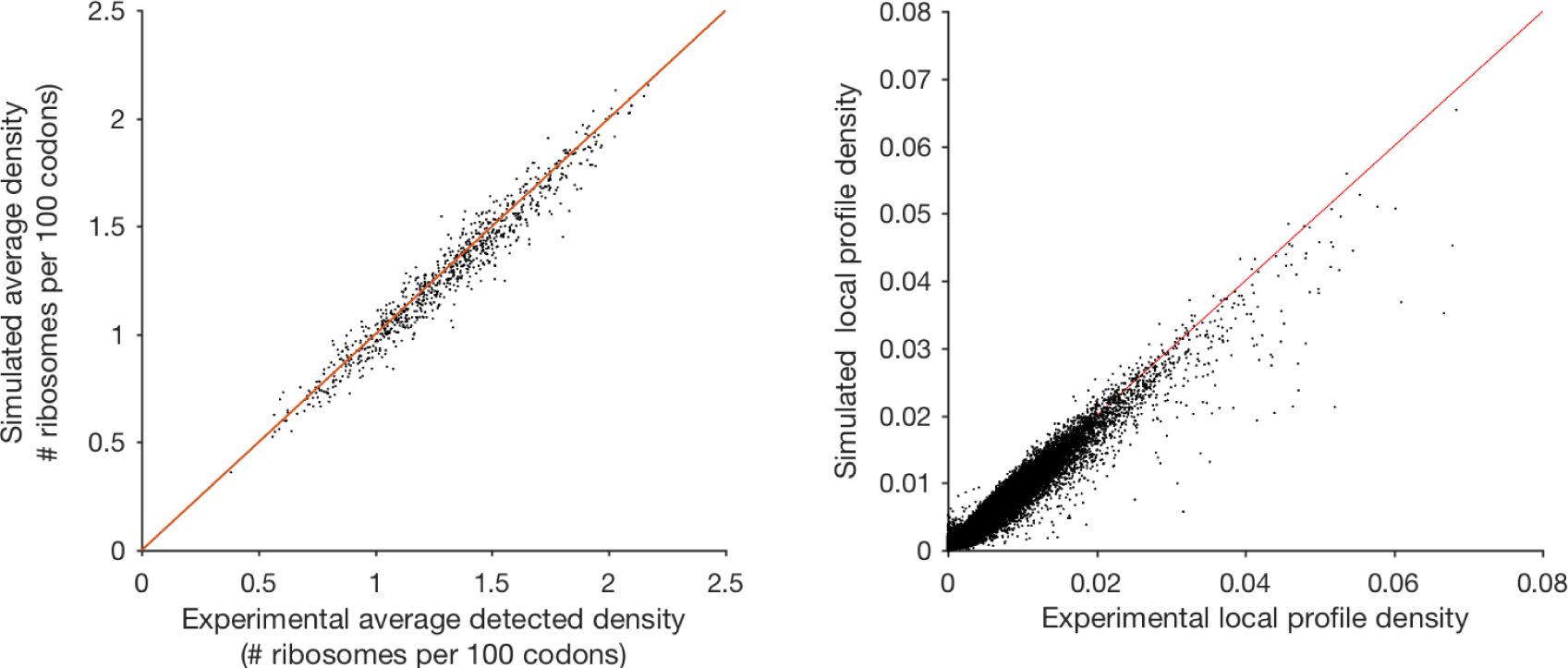
Comparison between experimental data and numerical results from inference procedure. We applied the inference method to a set of 850 genes in *S. Cerevisiae* (see **Materials and Methods**) and compared the total (left) and local (right) densities of the original dataset with the ones obtained by simulations of the model with the inferred parameters. The parameters used were 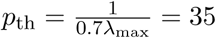 and λ_max_ = 50. The simulated and experimental densities were in good agreement, showing a Pearson’s correlation coefficient of 0.986 (p-value < 10^−5^). The individual profiles obtained by simulations also showed good agreement with the experiments, with Pearson’s correlation coefficient of 0.975 (p-value < 10^−5^).

**Figure S2.**
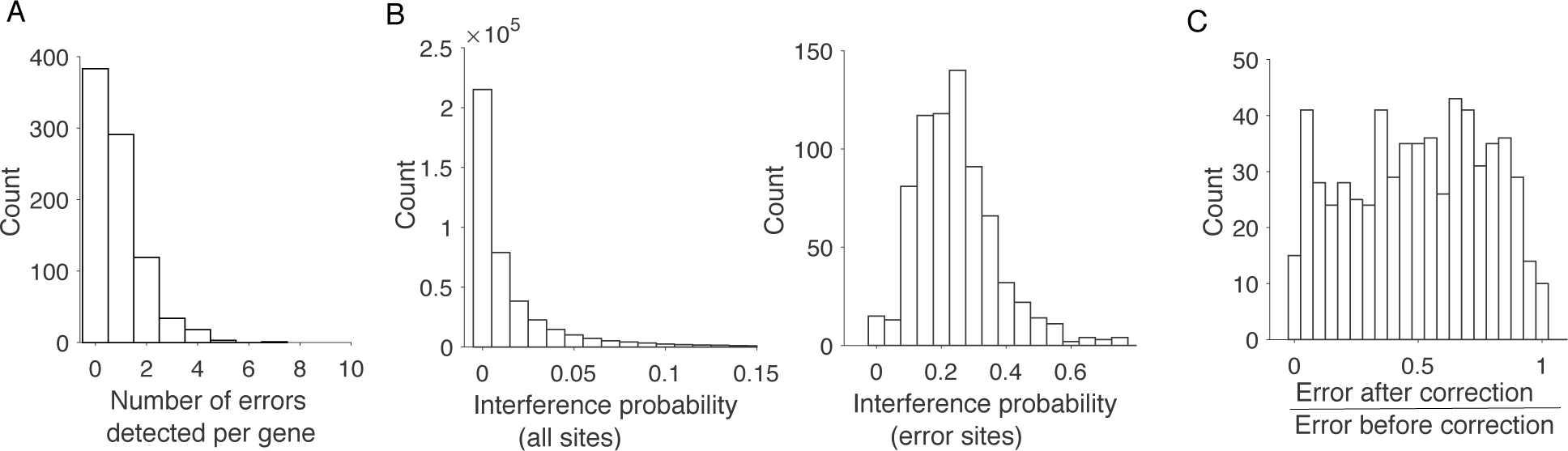
Detailed results of the inference procedure. **A**. Histogram of the number of significant errors detected for each gene. During the inference procedure, 383 genes over the 850 in the dataset (45%) did not require corrections, which means that the difference of profile observed between the simulation and the experiment was for these genes globally under the threshold error fixed by our procedure. For the remaining 467 genes, the number of error sites per gene above the threshold error was on average 1.57 (std = 0.925) **B**. Histogram of interference probability for significant error sites (left) and for all sites (right). We numerically estimated the probability of a ribosome occupying a certain site to block a ribosome located 10 codons before. We call this empirical probability the interference probability. We found that the interference probability of error sites was on average equal to 0.245 compared with an average rate of 0.011 over all the sites of our dataset. This large difference showed, as we expected, that local profile errors between experimental and simulations after the first round of estimation are primarily due to ribosomal interference. **C**. Histogram of error improvement after the refinement step, given by the ratio of error after correction over error before. The decrease in error was on average of 57%. For 65% of the sites the error after correction went under the initial threshold of error site detection. The reasons for correction failure can vary from too large initial error, configurations of error sites too close to allow separate correction and more generally possible missing reads or errors in the original data.

**Figure S3.**
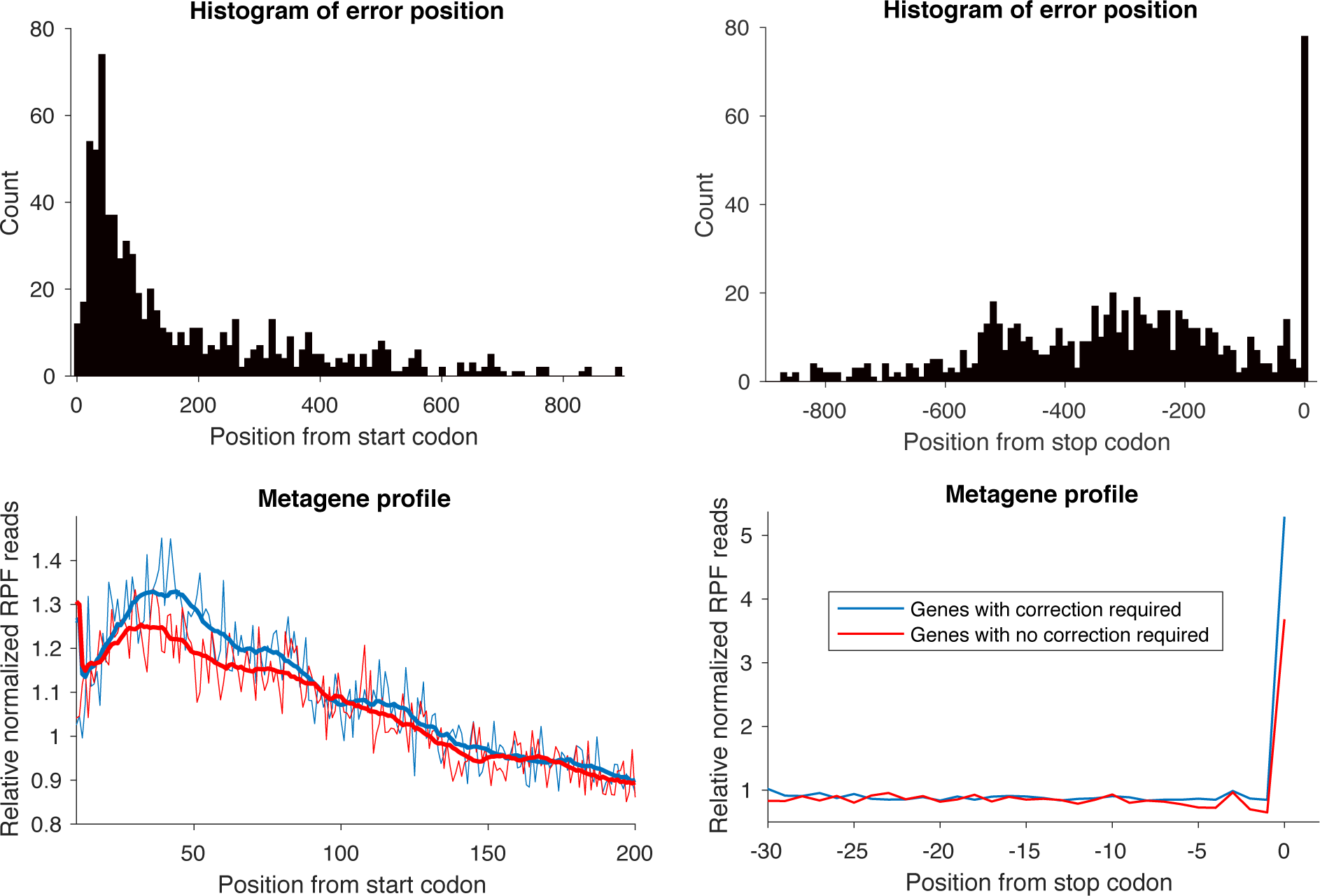
Positions and metagene profiles associated with correction. Up: Histograms of inconsistent sites obtained from running our procedure on our main dataset, after alignment from start (left) and end (right) position. **Down:** Comparison of metagene profiles (as in Ingolia *et al* [1]), after alignment from start (left) and end (right) position, between subset of genes that required correction procedure and genes that did not.

**Figure S4.**
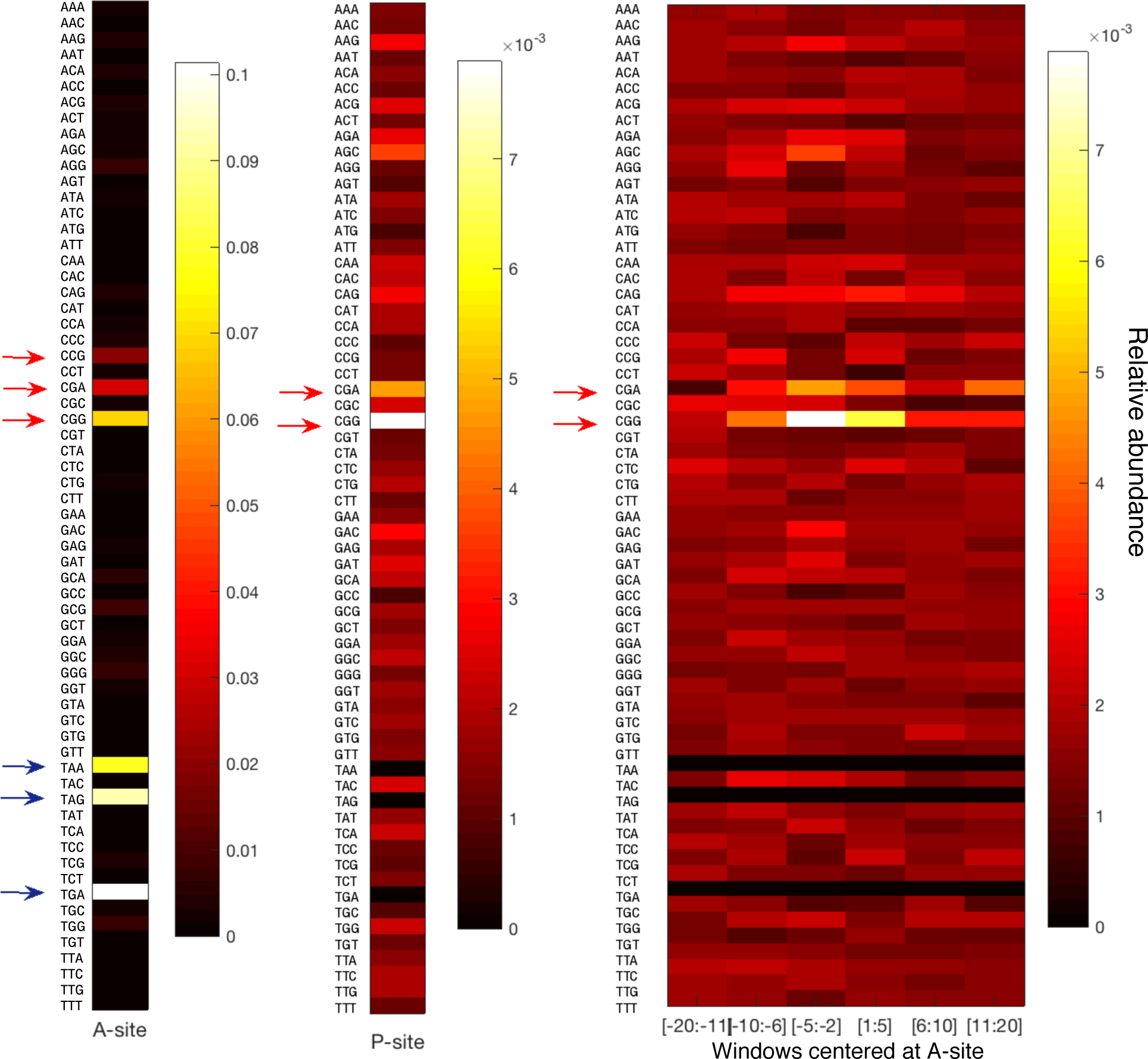
Analysis of codon context surrounding the inconsistent sites. The heatmaps show the relative abundance of each codon (ratio of the local frequency of the codon and its frequency over all the sites) at the A-site, P-site, and other windows ([−20:−11], [−10:−6],[−5:−2],[1:5],[6:10],[11:20]) at inconsistent sites. Blue arrows indicate stop codons. Red arrows indicate non stop codons with slowest mean elongation rate (see Figure 3).

**Figure S5.**
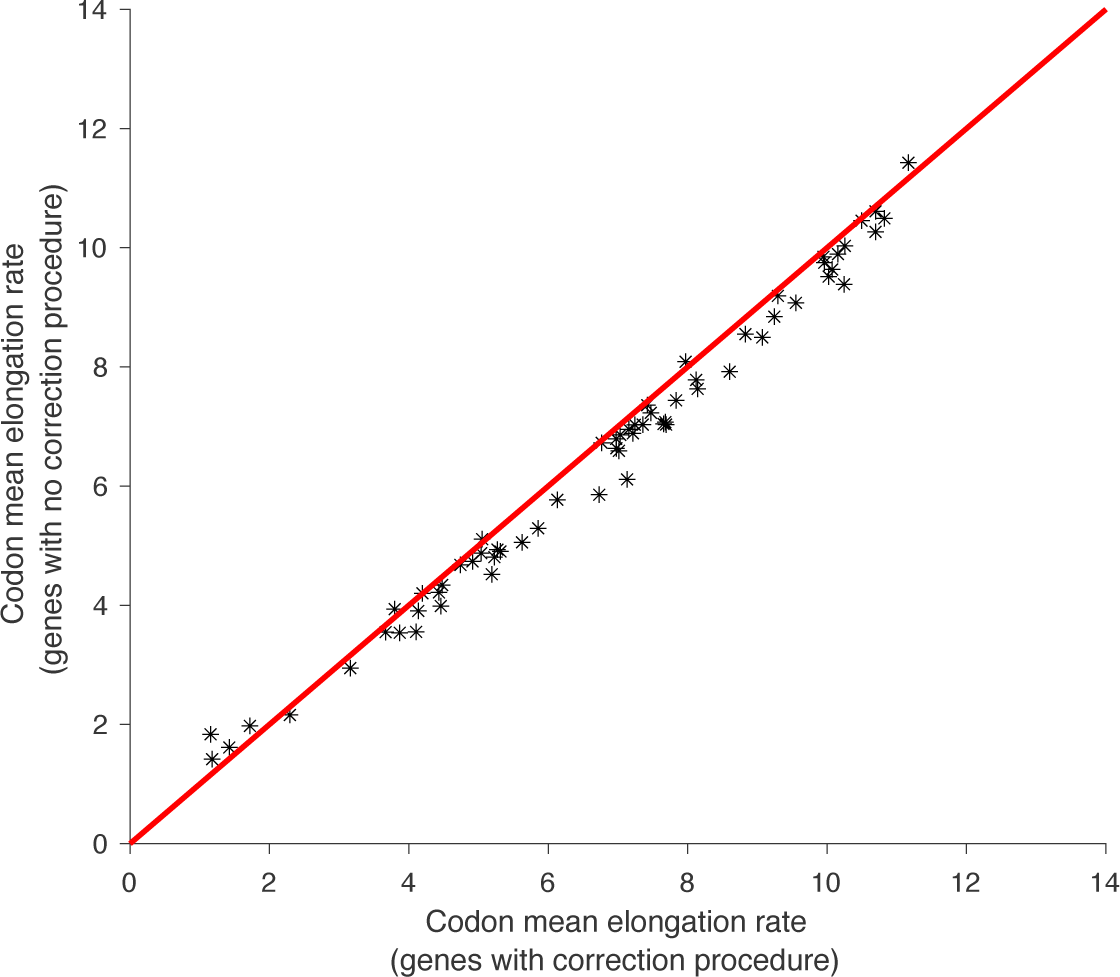
Comparison between codon MER of genes that required correction procedure and genes that did not.

**Figure S6.**
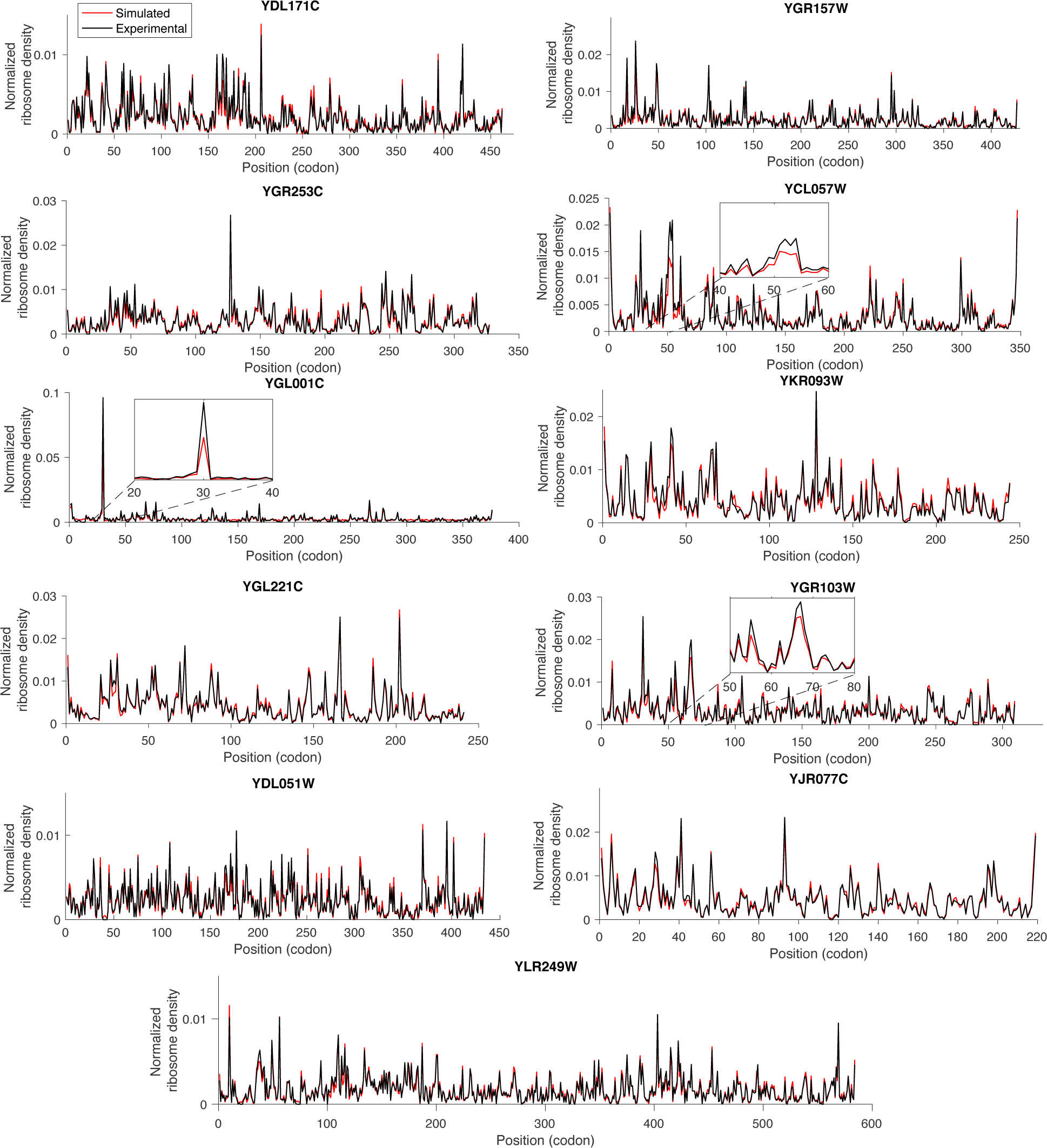
Examples of comparisons between experimental profiles (in black) and simulated ones (in red). The profiles are obtained by normalizing the ribosome density by the total number of reads. The inset panels (for genes YGR103W, YCL057W and YGL001C) show where the simulated model cannot reproduce the experimental profile. This notably happens when the average observed density is large enough so it gets incompatible with the presence of large peaks. In this case, the model cannot simulate a large peak density without having a queue which leads to stalled ribosomes, and the simulated density is lower than the observed one.

**Figure S7.**
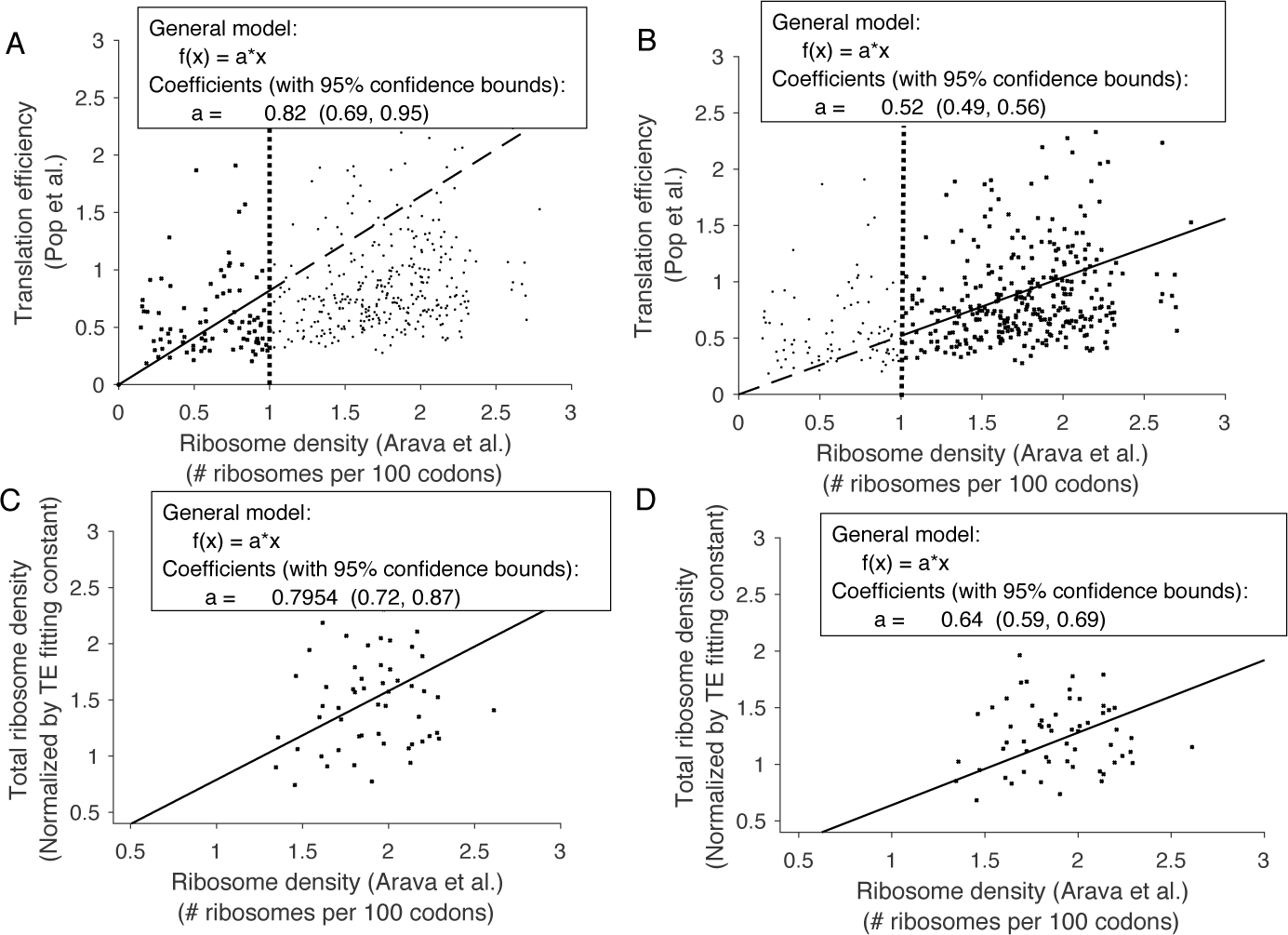
Comparison between translation efficiency (TE) from Pop *et al.* [19] and total ribosome density. Results from the linear fits are shown in inset. **A**. The gene-specific TE for 423 genes from Pop *et al.* [19] (see **Materials and Methods**) is plotted against the corresponding total ribosome density (average number of ribosomes per 100 codons) from Arava *et al.* [33]. We performed a linear fit of the points for which the corresponding ribosome density was less than 1 ribosome per 100 codons. **B**. Similar fit as in A in the range of ribosome density larger than 1 ribosome per 100 codons. **C**. The simulated total densities for a subset of 58 genes is plotted against the ribosome density from Arava *et al.* **D**. The simulated detected-ribosome densities for the same 58 genes is plotted against the ribosome density from Arava *et al.*

**Figure S8.**
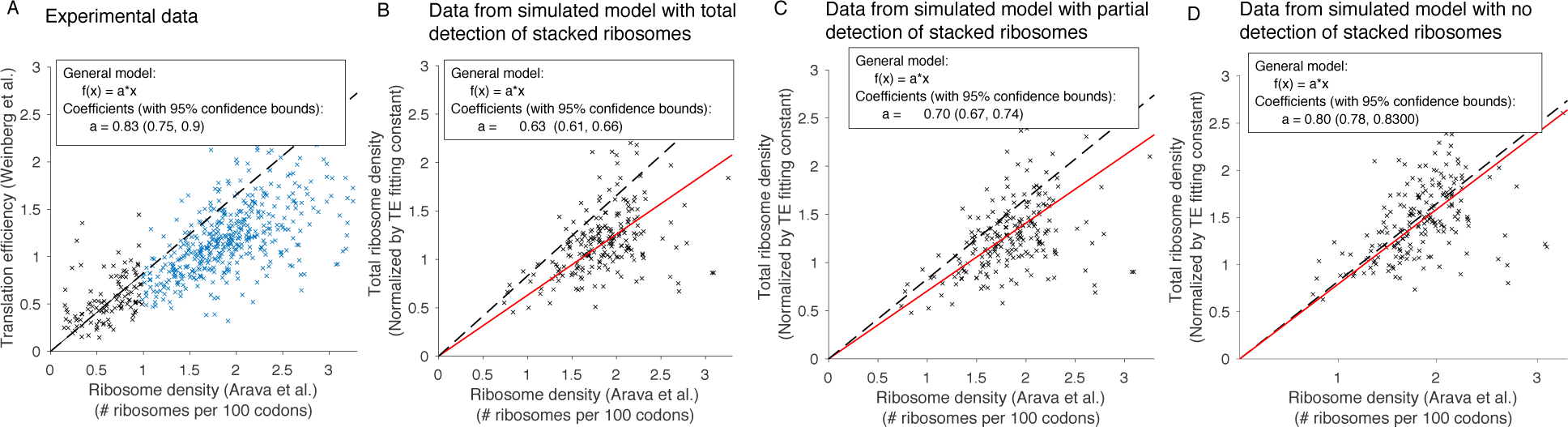
Comparison between ribosome average density measured in Arava *et al.* and translation efficiencies simulated under different profile models. **A**. We plot the experimental measurement of translation efficiency (ratio of ribosome RPKM and mRNA RPKM) and fit the TE to the density in the region where the density is less than 1 ribosome per 100 codons (linear coefficient 0.83). **B**. We compare the experimental density to the simulated TE under a model where all ribosomes get detected . The linear fit (plotted in red) between the simulated TE and the density gives a coefficient of 0.63. **C**. Same as in B, with a model where stacked ribosomes are partially detected with probability 0.5. Linear fit coefficient is 0.70. **D**. Same as B, with no detection of stacked ribosomes. Linear fit coefficient is 0.80, that is the best matching with experimental data .

**Figure S9.**
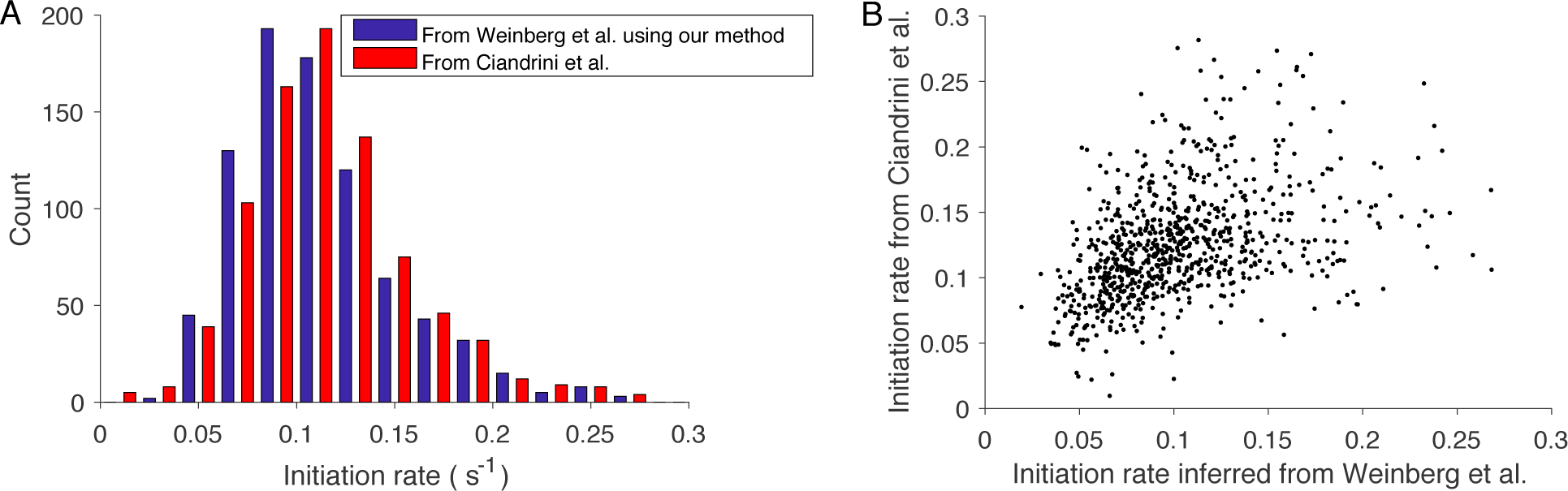
Comparison between the initiation rates inferred by applying our method to Weinberg *et al.* dataset and the ones inferred by Ciandrini *et al.* [34] from MacKay *et al.* [35]. A. Histogram of the two sets of inferred initiation rates. **B**. Comparison between initiation rates inferred by the two methods. Pearson *R* = 0.4, p-value< 10^−5^.

**Figure S10.**
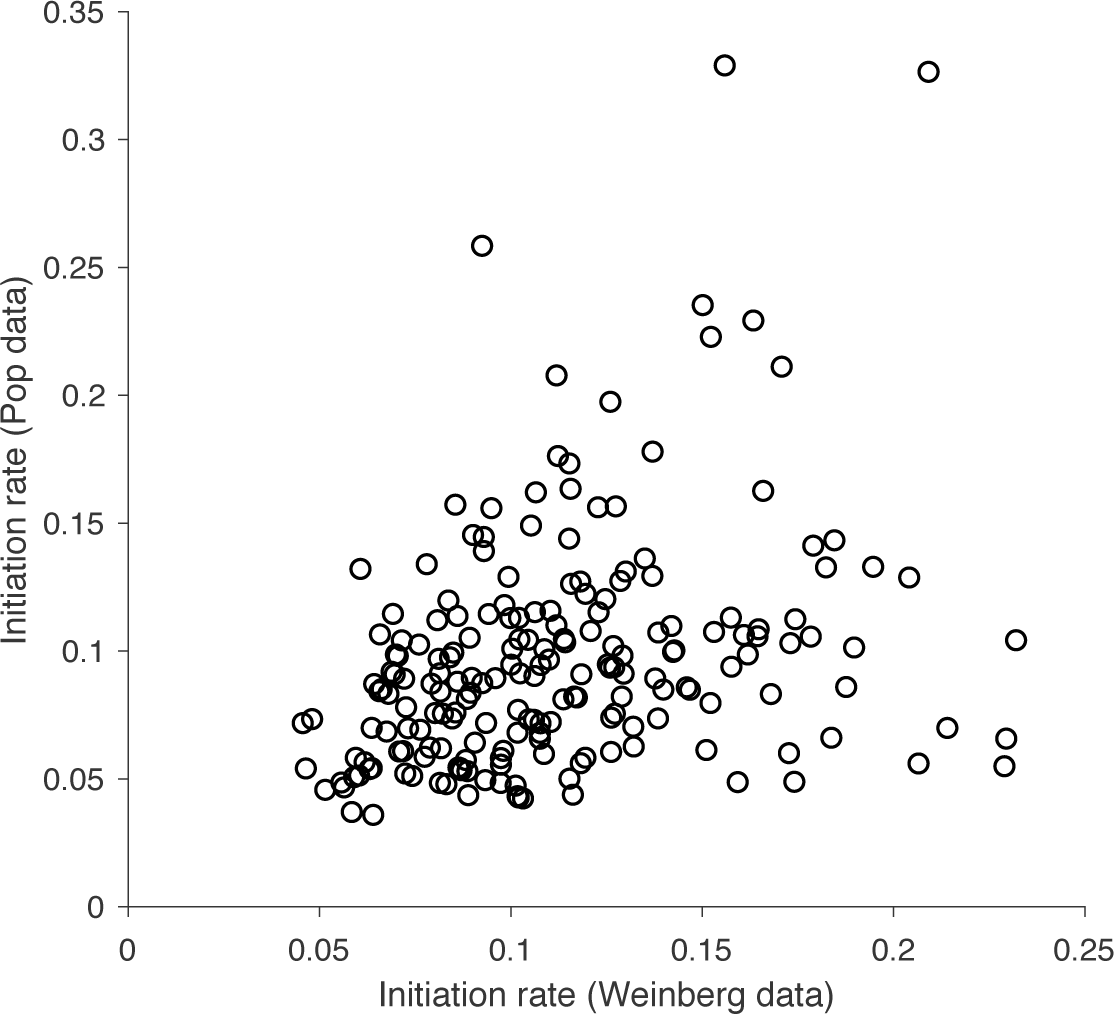
Comparison between the initiation rates inferred by applying our method to Weinberg *et al.* dataset [16] and to Pop *et al.* [19] (212 genes in common). Pearson *R* = 0.31, p-value< 10^−5^.

**Figure S11.**
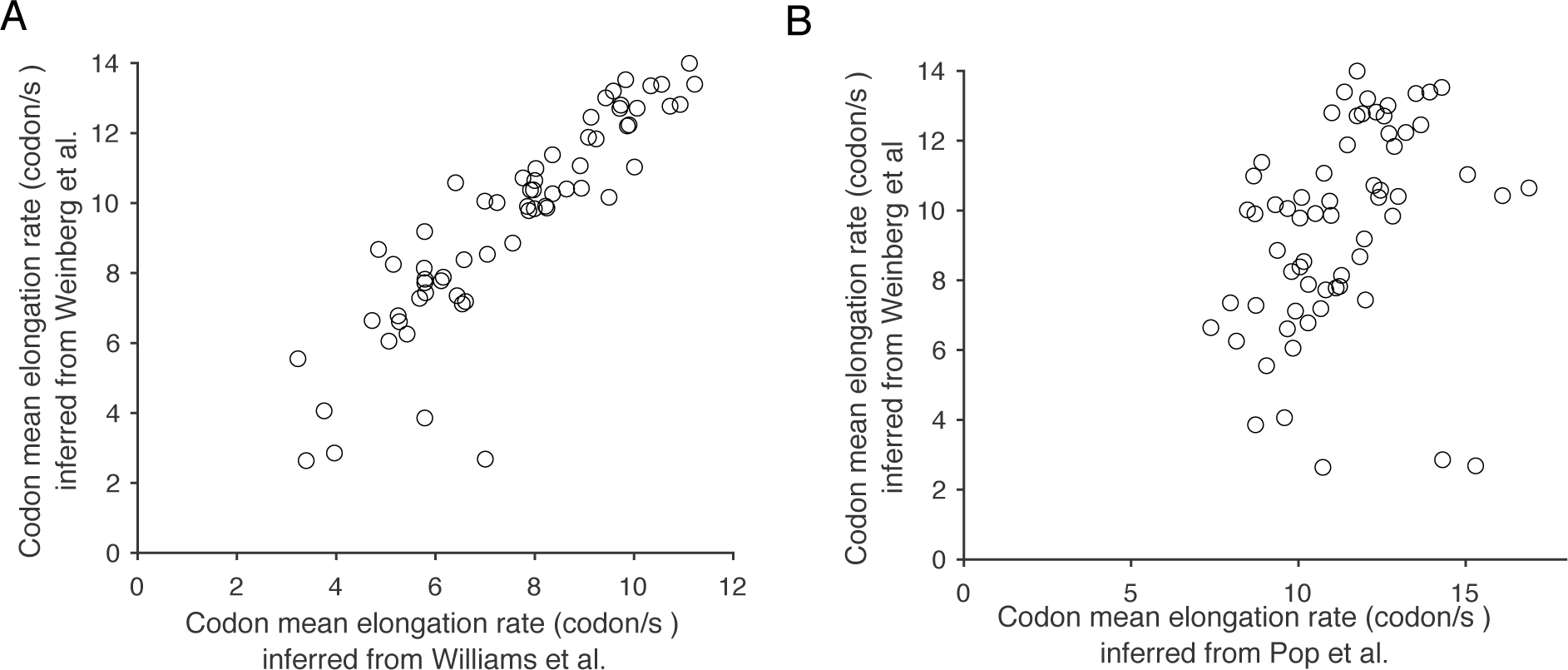
Comparison between codon-specific mean elongation rates. **A**. We compare the codon-specific mean elongation rates inferred from the Weinberg *et al.* dataset [16] to the ones inferred from Williams *et al.* dataset [32]. **B** Comparison between the codon-specific mean elongation rates inferred from the Weinberg *et al.* dataset [16] and the ones inferred from Pop *et al.* dataset [19].

**Figure S12.**
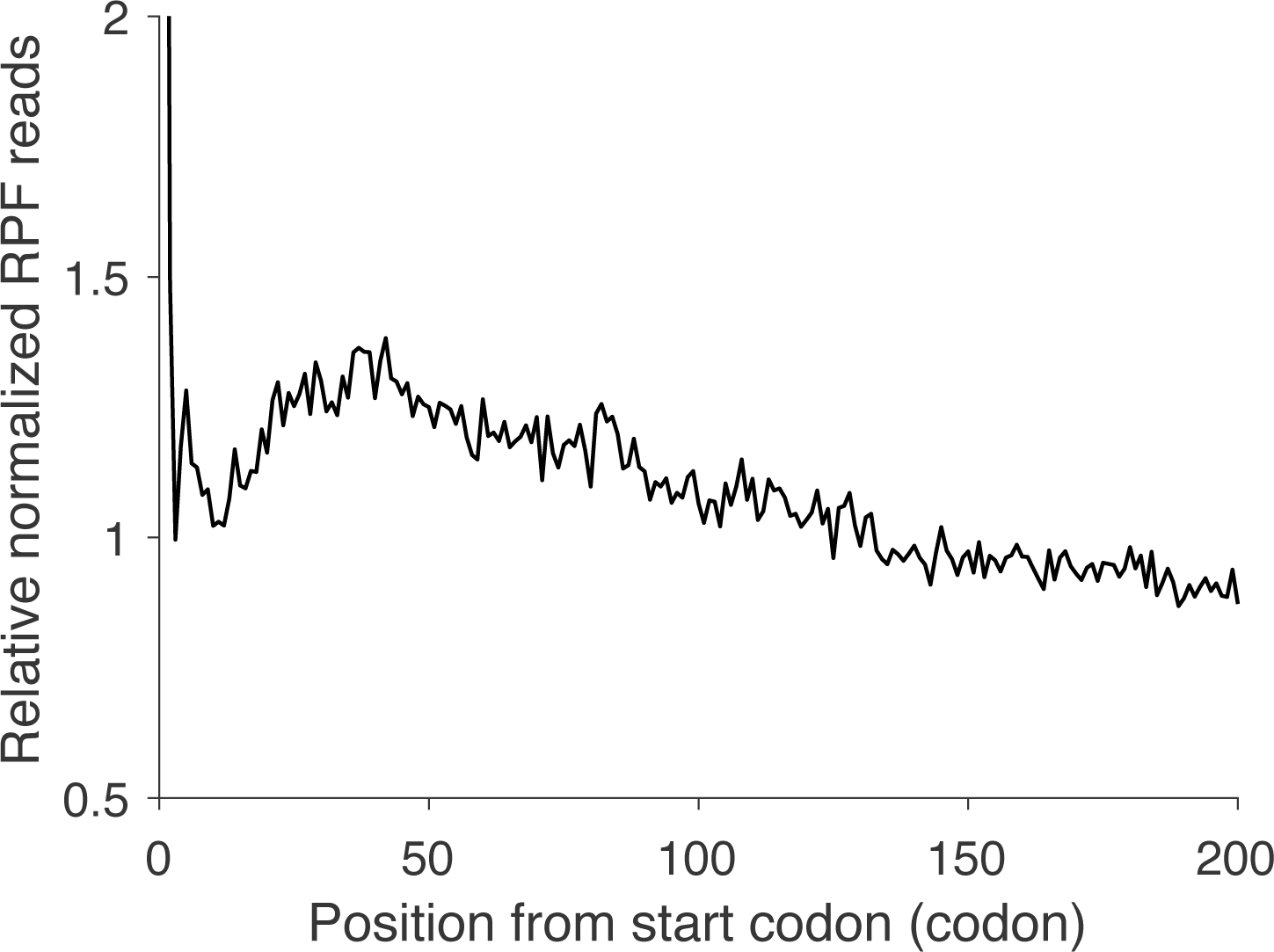
Metagene relative normalized ribosome-footprint density as a function of codon position.Ribosome profile footprint (RPF) reads in open reading frames (ORFs) from Weinberg *et al.* [16] were individually normalized by the mean RPF reads within the ORF, aligned from start codon and then averaged with equal weight for each codon position across all ORFs, as in Ingolia *et al.* [1].

**Figure S13.**
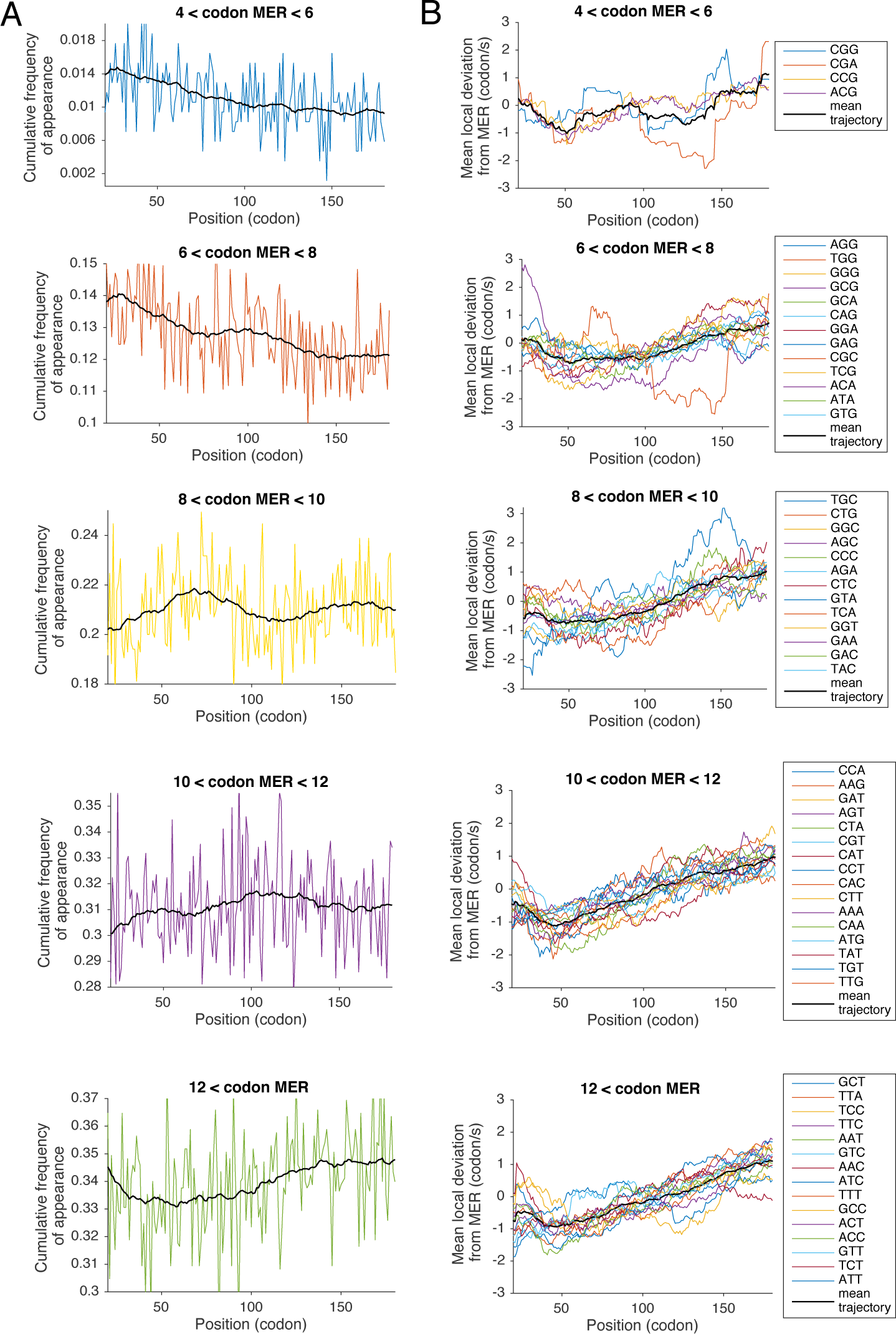
Detailed codon frequency of appearance and elongation speed along the transcript. **A**. Different panels show the frequency of appearance for each group of codons from Figure 5A. The black curve in each panel corresponds to a smoothed version, for which the value at position *i* is obtained by averaging the values between positions *i —* 20 and *i* + 20. B. The difference between codon-specific local speed shown in Figure 5B and the average of codon-specific speeds between position 20 and 180. Different codons are grouped as in A. For each panel, the black curve corresponds to an average of the curves in that panel.

**Figure S14.**
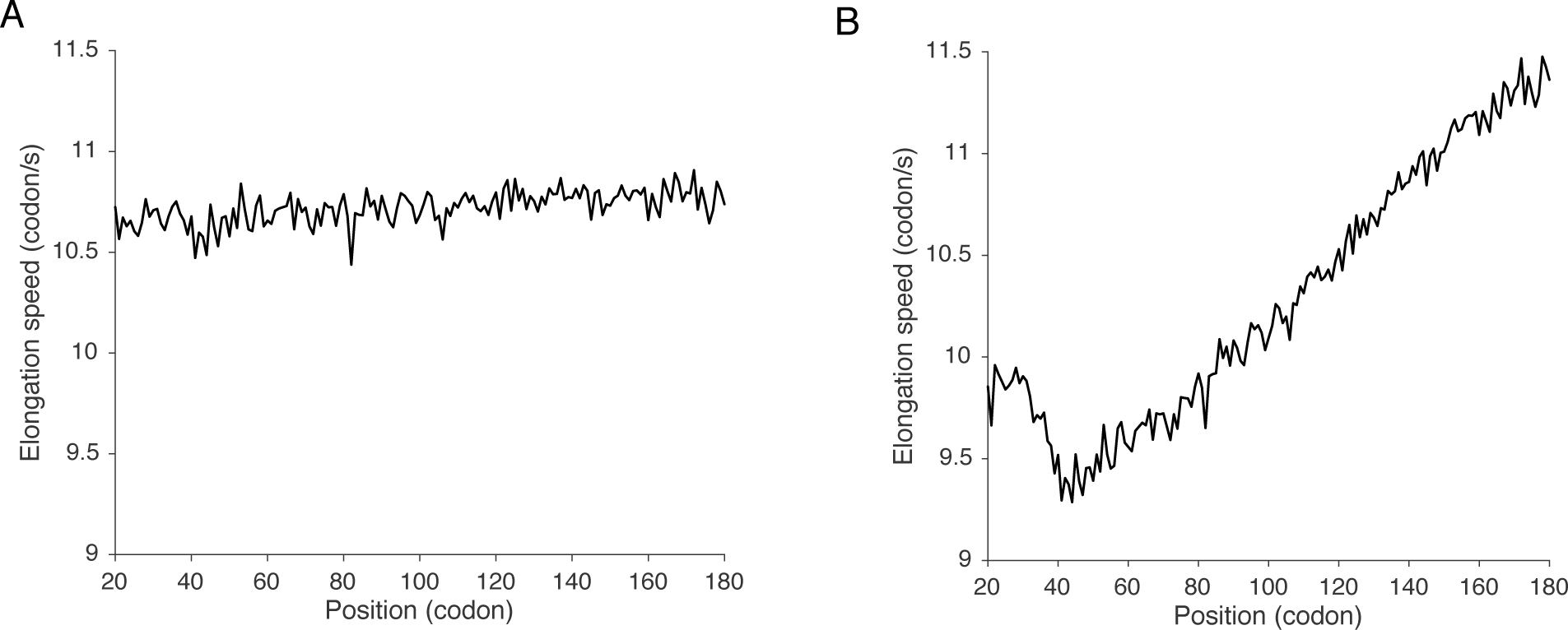
A. Average elongation speed along the transcript obtained by setting the elongation speed for each codon type at all positions to the corresponding mean elongation speed computed from Figure 3B. This plot shows that the variation of codon frequency along the transcript is not sufficient to explain the 5′ translational ramp. **B.** Average elongation speed along the transcript obtained by setting the elongation speed for each codon to the position-specific mean elongation rate in Figure 5B.

**Figure S15.**
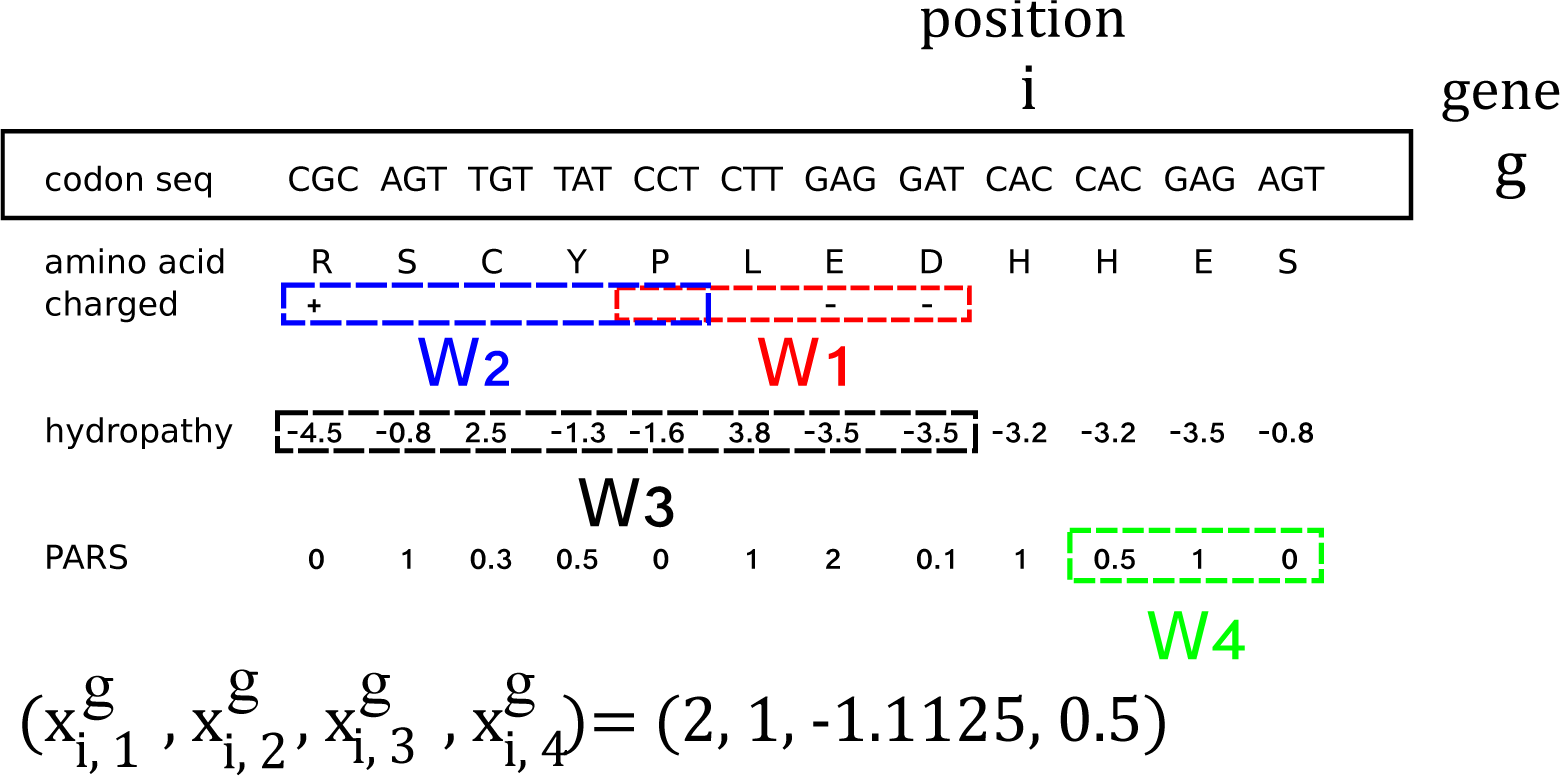
An example of variables used to fit the deviation to codon mean elongation rate (equation (1)). At position *i*, we consider 4 types of variables in our fitting models: *x_i,1_* is associated with the negative charges, *X_i,2_* with the positive ones, *x_i,3_* with the hydropathy score and *x_i,4_* with the PARS score. For each variable *x_i,1_*, a window *W_k_* is defined, such that 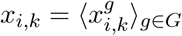, where *G* is the gene set, and for a gene *g*,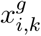 is computed by averaging (for hydropathy and PARS score) or counting (for the number of charges) the associated feature over positions *i — W_k_* for the charges and hydropathy scores, and positions *i* + *W_k_* for the PARS score. In this example, *W_1_* = [1:4], *W_2_* = [4:8], *W_3_* = [1 : 8] and *W_4_* = [1:3].

**Figure S16.**
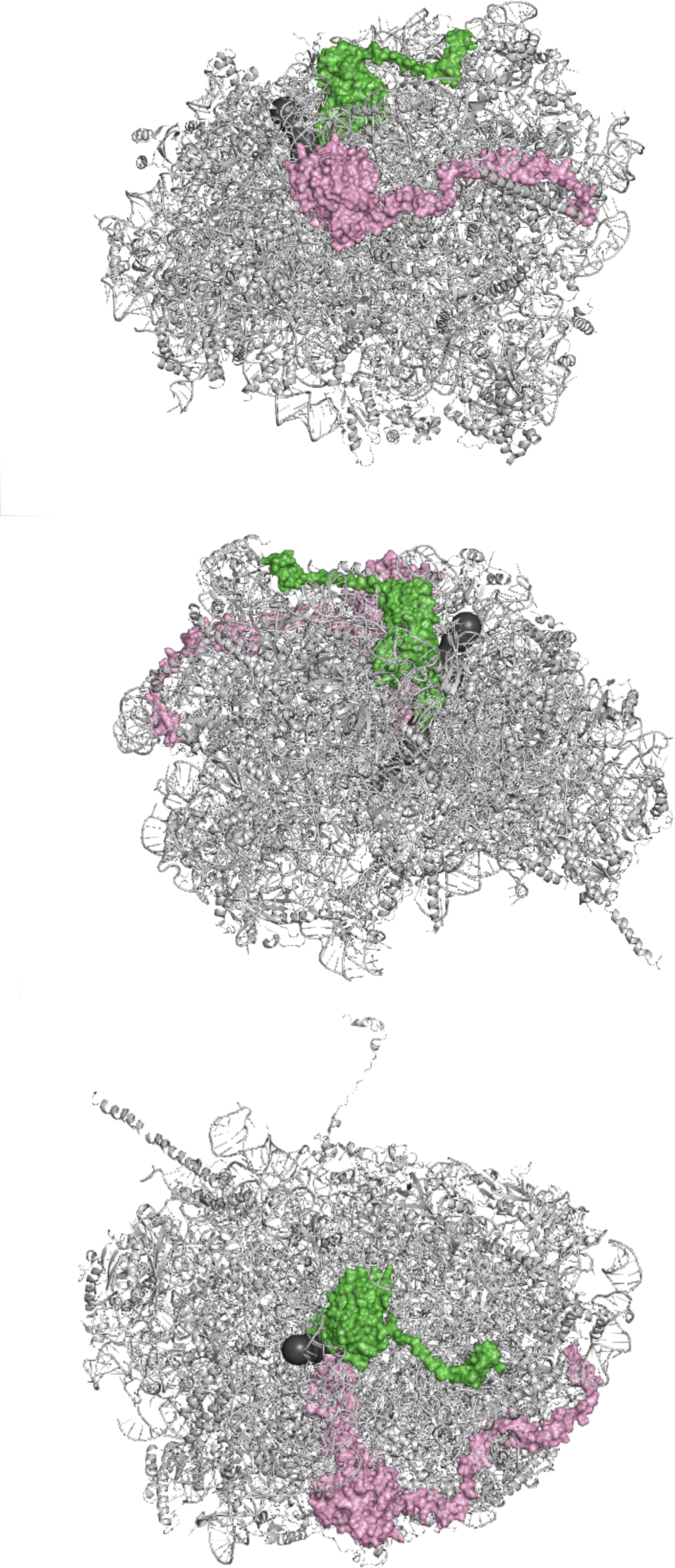
The atomic structure of the large ribosomal subunit in yeast (under three different viewing angles), with the ribosome exit tunnel (in black) and proteins L4 and L22 (in pink and green, respectively).

**Figure S17.**
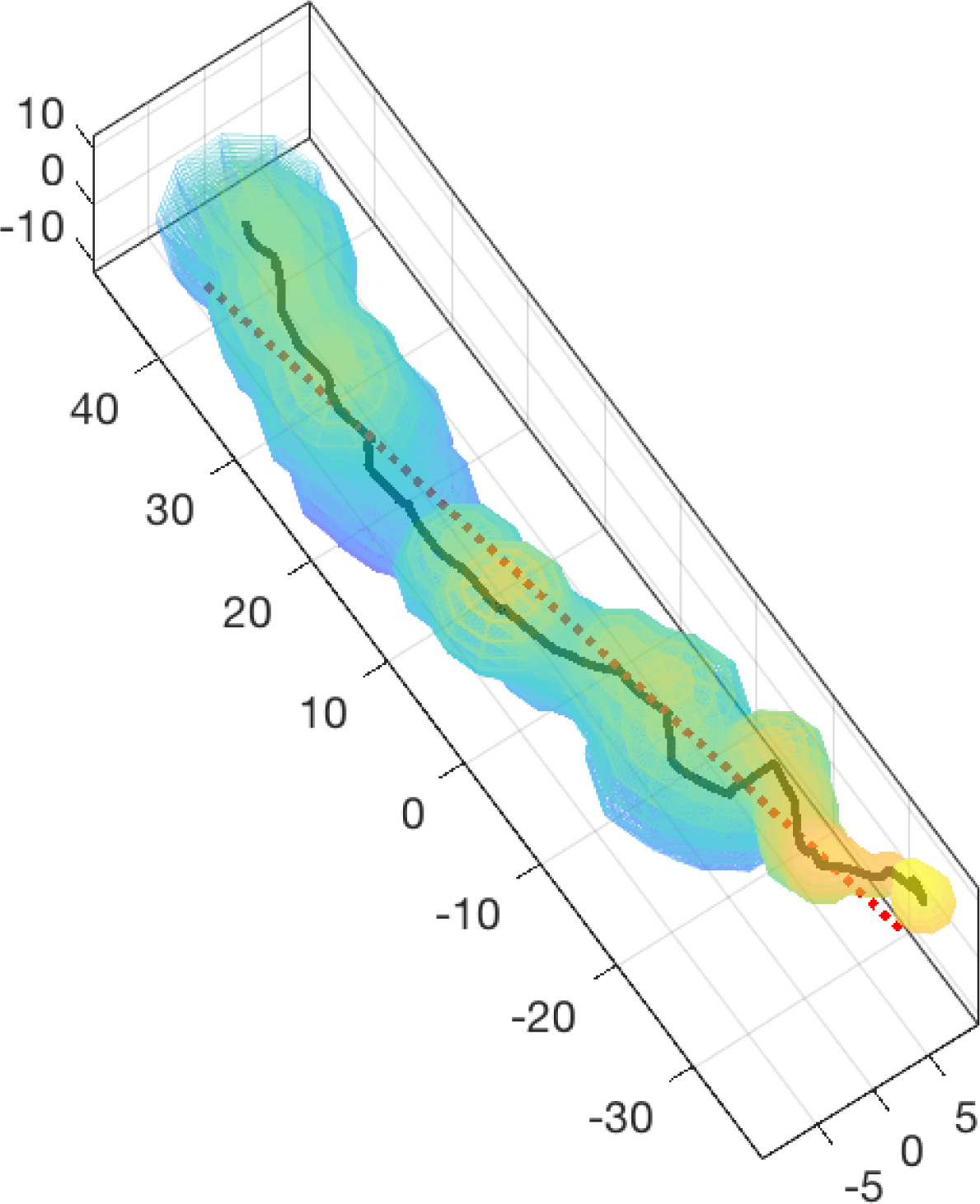
A detailed view of the ribosome exit tunnel extracted from cryo-EM data. We plot (in black) the centerline of the tunnel (see **Material and Methods**) from the PTC (bottom right) to the exit (top left)). A linear fit of the centerline (coefficient of determination *R*^2^ = 0.985) is shown in red.

**Figure S18.**
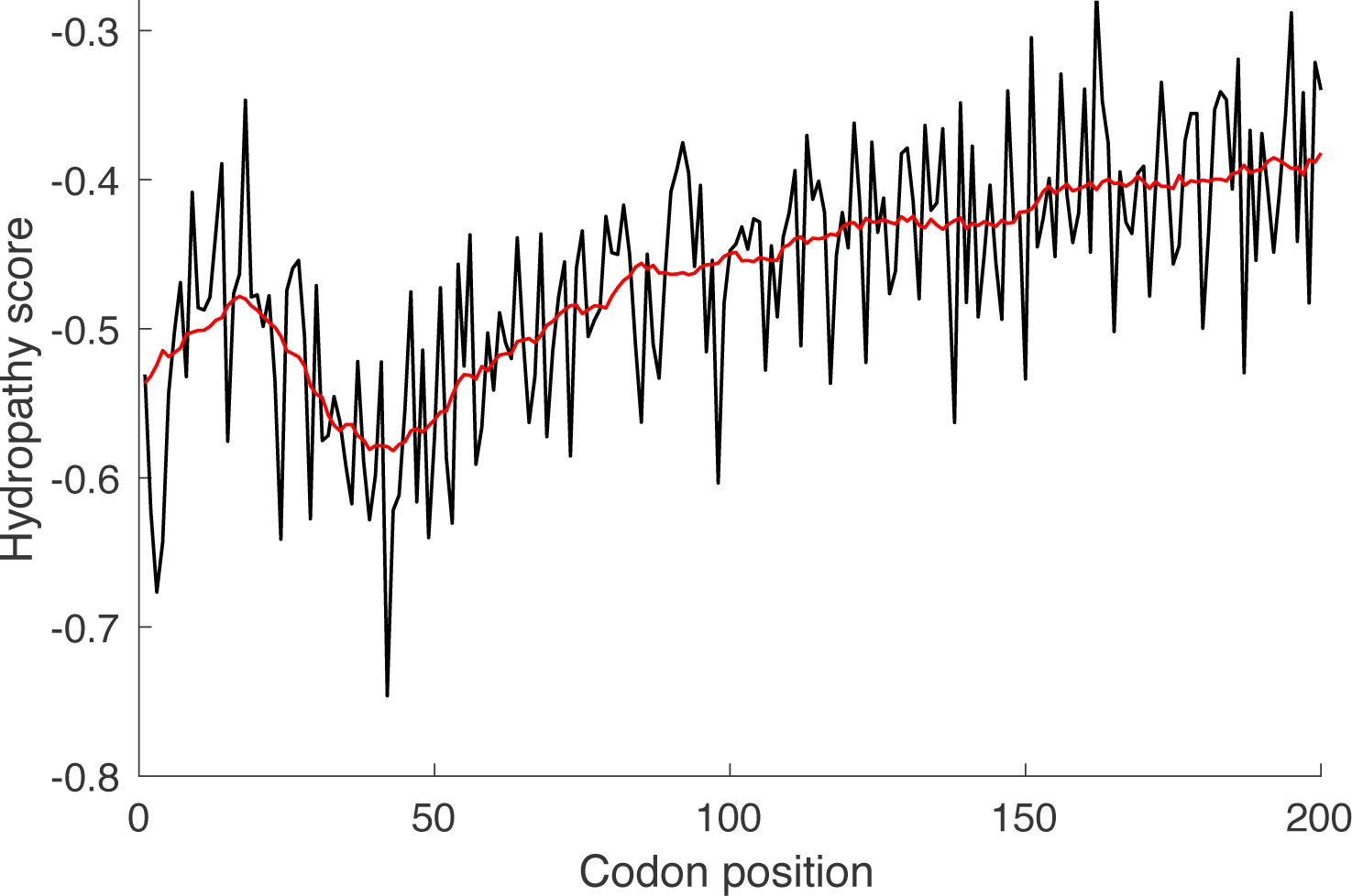
The average hydropathy score across codon positions. We plot (in black) along the codon position the hydropathy scores averaged over all genes of the non-filtered dataset (2862 genes). In red, we plot this average score smoothed by averaging over a 10 codons window.

**Figure S19.**
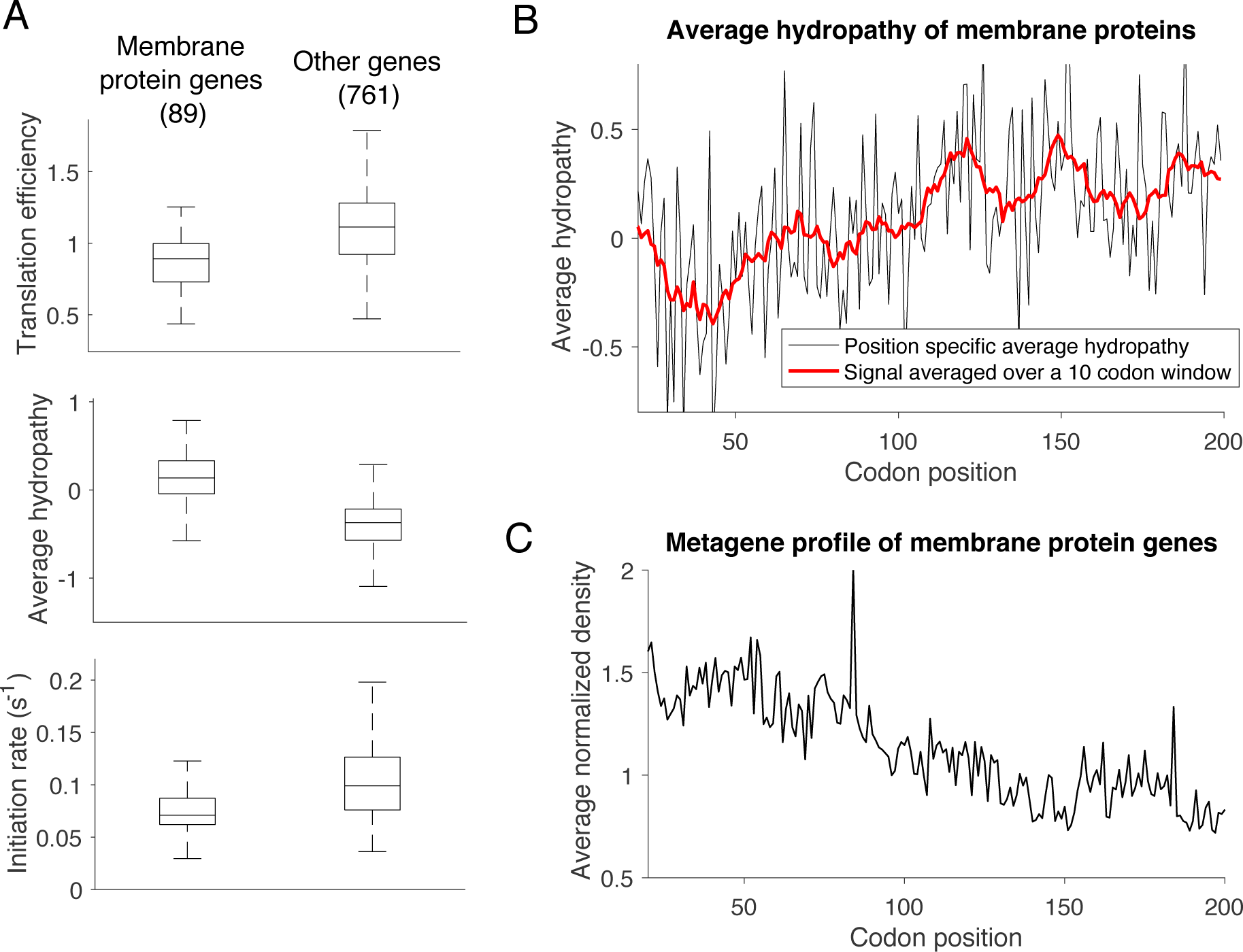
Analysis of membrane protein genes. **A**. We compare the translation efficiency, average hydropathy and the inferred initiation rates of a subset of 89 membrane protein genes to the other genes of our main dataset. These 89 genes were obtained by cross-referencing our main list of genes (850 genes) to a list of 666 genes associated with membrane proteins (from Miller *et. al.* [55], table 2). We found that the membrane protein genes have in average lower TE (0.88 ± 0.20 compared to 1.10 ± 0.26), larger hydropathy score (0.14 ± 0.28 compared to —0.42 ± 0.3) and lower initiation rates (0.07 ± 0.02 *s*^−1^ compared to 0.11 ± 0.05 *s*^−1^). Boxplots give the lower and upper adjacent values, first and third quartile and median values. **B**. The average hydropathy score across codon positions for membrane protein genes. We plot (in black) along the codon position the hydropathy scores averaged over all 89 genes. In red, we plot this average score smoothed by averaging over a 10 codons window (see also Figure S18). **C**. Metagene relative normalized ribosome-footprint density as a function of codon position for the 89 membrane protein genes (see also Figure S12).

**Figure S20.**
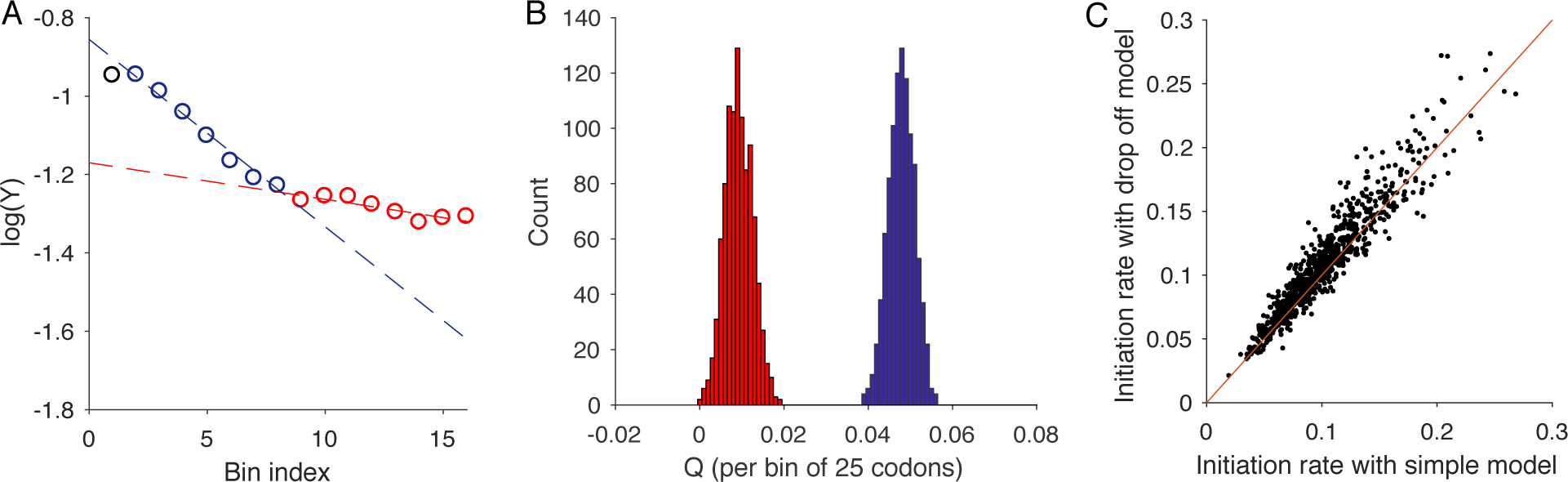
Estimation of drop off rate using Sin *et* al.’s methods [22]. **A**. We divide the ORF in bins of 25 codons and plot for each bin the logarithm of the average normalized number of reads counted from Weinberg *et al.* dataset [16]. The associated drop off rate is estimated by linearly fitting the average value of each column with *Ae*^-QX^, where *X* is the bin index and *A, Q* are fitting parameters. We distinguish two regions of constant drop off rates in windows [25, 225] (in blue) and [200,400] (in red). Linear fitting (dotted lines) gave *Q* = 0.048, with resulting drop-off probability per codon and per elongation event *r* = 0.002) for region [25,225] and *Q* = 0.0093 (*r* = 3.7.10^-4^) for region [200, 400]. Genes were chosen to be of length > 400 codons and such that the number of reads per codon is on average larger than 10 (458 genes). **B**. Histogram of *Q* following the bootstrap method from Sin *et al.* [22] (number of samples 10^3^). C. We compare the original estimates of initiation rates with the ones obtained with a modified model of translation including drop-off with probability 3.7 × 10^4^ (Pearson *R* = 0.94, p-value < 10^−5^).

**Figure S21.**
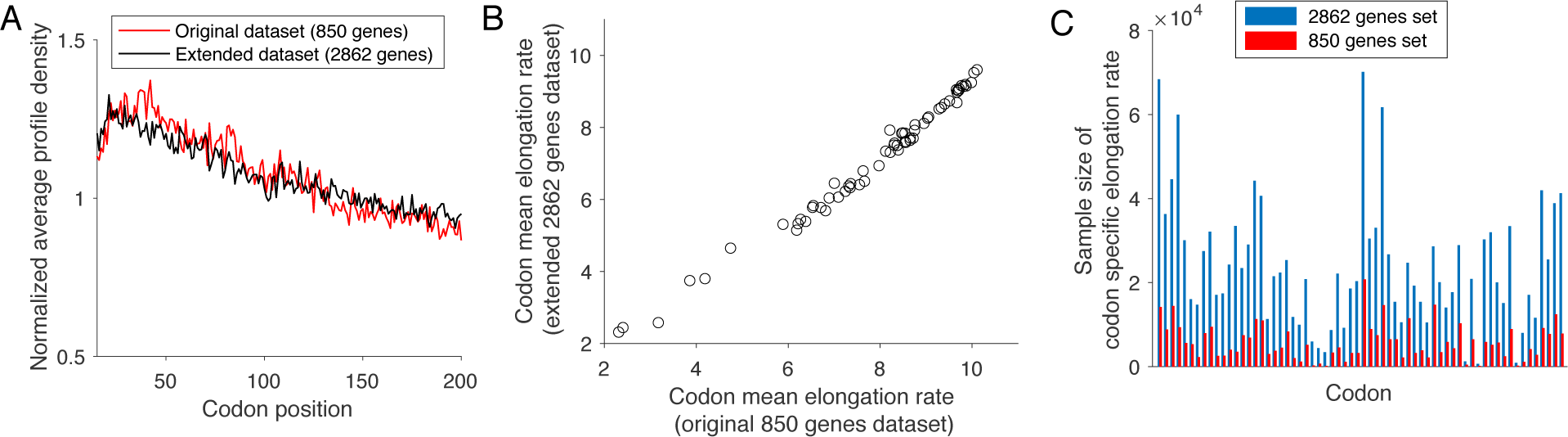
Comparison between results on the dataset mainly used in our analysis (850 genes, see Methods section) and the extended one (2862 genes) shows no bias due to filtering. **A**. We plot (in black) the normalized average profile density across the first 200 codons (as done in Figure S12) from the extended dataset of 2862 genes (selected to be of length > 200 codons). We compare these variations to the ones obtained (in red) from the filtered dataset used in our main analysis (length > 200 codons and average number of footprints > 10 per codon). Both show the same 5′ “translational ramp” pattern. To obtain the variations for the extended dataset at each position *i*, the genes whose profile at *i* contained at least one footprint were considered (for any gene *g*, *p_g_*(*i*) > 0). These profile values were averaged across the genes, after normalizing each of them by *Σ_k∊I_g__ p_g_* (*k*), where *I_g_* is the set of all positions *k* where *p_g_* (*k*) > 0 (such processing was done to discard the gene-specific positions that contains no information on the elongation rate variations). **B**. Comparison between the mean codon elongation rate from the filtered and non-filtered dataset. The elongation rates were obtained using the naive estimates of our inference procedure (see Supplementary section) and filtered by selecting rates < 20 codon/*s*. This filtering was done to discard the estimates from positions with no or a few footprints, as these estimates are not informative of the dynamics (the threshold of 20 codon/*s* still allows to keep more than 85 % of the inferred rates, for the codon with largest mean elongation rate). **C**. We plot for each codon and each dataset the sample size of the associated elongation rates used to get the mean codon elongation rate in (B). This comparison shows that after filtering for rates < 20 codon/*s*, the extended dataset of 2862 genes still contains for each codon on average ~ 5.3 more rates than the original dataset of 850 genes.

